# Entangled stoichiometric objectives shape microbial catabolism

**DOI:** 10.1101/2025.07.29.667445

**Authors:** Maaike Remeijer, Frank Bruggeman

## Abstract

The search for fundamental relationships between energetic and biosynthetic parameters of catabolism and anabolism is a major goal in microbiology. This is complicated by the fact that ATP synthesis is required for some anabolic precursors, all building blocks, and their polymerization into macromolecules, while the synthesis of other anabolic precursors and catabolic products yields ATP. Yield parameters were classically predicted from approximate phenomenological relations between catabolic and anabolic stoichiometry. Here we compare the catabolisms of a diverse set of microbial species across conditions using genome-scale stoichiometric models. We focus on states of maximal energetic efficiency (maximal yield of biomass of the energy source) and present an unbiased method for calculating stoichiometric relations between catabolism and anabolism. We find that synthesis of charged energy-carriers and anabolic precursors by catabolism is strongly intertwined. Catabolic intermediates and reactions vary greatly, due to variation in the energy and carbon source for growth. We find that the ATP requirement for 1 gram biomass varies between 72.8 and 246.1 moles, precursor sets vary between 4 and 14 in size, and acetyl-CoA is the only common precursor across species. We conclude that the complex interplay between precursor synthesis and energy conservation of heterotrophic catabolism results from an optimal compromise between conflicting objectives. The state of maximal energetic efficiency is reached by minimizing the carbon source lost during energy catabolism due to catabolic-product formation. This choice is influenced by the need for an optimal precursor set that compromises between maximal ATP production during its formation from the carbon source and minimal ATP consumption when it is converted into building blocks. We find that the associated optimal catabolic pathways are diverse across species and conditions.

## 1 Introduction

The plasticity of catabolism largely explains the omnipresence of microbes. Catabolism is an adaptation to the available sources of energy and chemical elements while biosynthesis (anabolism) is condition invariant and conserved. Neidhardt, Ingraham and Schaechter [1] expressed this invariance as “*The chemistry of biosynthesis has one marvelously simple aspect: virtually all biosynthetic pathways, polymerization and assembly processes are fundamentally the same in all bacteria*.” [1]. They also suggested that “*Regardless of which fueling reactions an organism employs, it synthesizes the same 12 precursor metabolites; they are the metabolic link between fueling and biosynthesis*.”. These concepts motivated [1] them and others [2, 3, 4, 5, 6, 7, 8, 9, 10, 11] to search for a rational and principled approach to the stoichiometry of microbial metabolism. In this paper, we will revisit these concepts and assertions using genome-scale stoichiometric models of metabolism (GSMMs) of different microbes to arrive at a more unbiased analysis, based on reaction stoichiometry and less on heuristics.

Neidhardt, Ingraham and Schaechter [1] distinguished four metabolic processes that are also central to this work. Here we list them for heterotrophic growth:

1. ‘fueling pathways’ convert the carbon and energy source (CS) into a catabolic product (CP; e.g. carbon dioxide, fermentation products), charged energy equivalents (EE; ATP, NAD(P)H) and precursor metabolites (PM),
2. ‘biosynthetic pathways’ convert the precursor metabolites into building block metabolites (BB; i.e. nucleic acids, amino acids, fatty acids, etc.), consuming charged energy equivalents and additional chemical element sources from the environment.
3. ‘polymerisation’ produces macromolecules (MM; e.g. protein, D/RNA, lipopoysaccharides, storage molecules) from the building blocks, consuming charged energy equivalents,
4. ‘assemblage’ achieves the macromolecular make up of 1 gram cells and consumes charged energy equivalents for maintenance processes.

We shall refer to process 1 as catabolism and 2-4 as anabolism. We aim to identify these processes and their net stoichoimetry in computed metabolic-states of GSMMs as function of the energy source.

Bauchop and Elsden [12] found that the grams of biomass (its dry weight) obtained from 1 mol of energy source (the biomass yield on energy source) varied more with growth conditions than the biomass yield on ATP (*Y*_*X/ATP*_). This, they argued, was due to the variable ATP yield of fueling pathways on their respective energy sources and to the invariance of anabolism. They, and others [9], postulated that *Y*_*X/ATP*_ was a biological constant (equal to about 10 gram drw/mol ATP); since this is an anabolic parameter, independent of the fueling pathway. Stouthamer and Bettenhausen (1973) [13] argued that it is a biological constant only when corrected for the maintenance requirement [14]. This corrected parameter was called 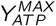. Stouthamer [2, 3] subsequently calculated its value across several conditions and found that it varied. Why its value varied, how good his estimates are, and why his aerobic glucose estimate was nearly a factor of 2 too large (called the ‘*Y*_*ATP*_ problem’ [15, 16]) has so far not been addressed. In this paper, we revisit these questions and relate them to the two functions of catabolism: synthesis of charged energy equivalents and precursor metabolites. We will show that it is their intertwined nature that makes *Y*_*X/ATP*_ (or 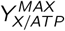) so hard to predict and vary with conditions.

A limitation of the calculations by Stouthamer [2, 3], Forrest & Walker [9] and Umbarger [4] is that a steady-state flux distribution is assumed, but not verified to be correct. This shortcoming was overcome by the mathematical approach called flux balance analysis (FBA) [17], initially proposed by Varma & Palsson [18, 19, 6]. They constructed stoichiometric models of metabolism and computed steady-state flux vectors that satisfy steady-state and flux-bound constraints and subsequently maximized a metabolic objective such as biomass yield on the energy source. This approach led to the GSMMs that are widely used now [20] and reconstructed from sequenced genomes [21]. These models exploit the environmental independency of the polymerisation and assemblage processes mentioned above by subsuming them all in a single reaction – called the ‘biomass reaction’ [17] – that converts all the building blocks together with an ATP cost (polymerisation and maintenance) into 1 gram biomass. Here, we will use FBA to calculate metabolic states of optimal energetic efficiency, the associated precursor sets and energy requirements for different microbial species and conditions.

GSMMs did for a long time not exist for non-model microbes studied in microbial ecology and biotechnology. (This situation is rapidly changing now with automatic GSMM reconstruction tools from genome information alone [22, 23].) These disciplines were therefore forced to use approximate methods to estimate biomass and catabolic-product yields on energy source [5, 10, 11, 8]. *Y*_*X/ATP*_ is a key parameter in these estimates. Since it equals the anabolic demand of ATP, it specifies the activity of catabolism needed for biosynthesis of 1 gram biomass. Given the ATP yield on the energy source by catabolism, the biomass and catabolic-product yield on energy source can be calculated. These calculations are based on net stoichiometries of catabolism and anabolism (called macrochemical equations [10, 24, 8], whereas we use GSMMs to obtain exact values of the bioenergetic parameters.

Many of the studies mentioned above recognized that precursor biosynthesis and charging of energy equivalents are intimately linked in fueling pathways and dependent on the available energy source. They realized that ATP synthesis may occur during precursor synthesis. The outcome of these calculations therefore depend on the assumed precursor set and associated synthesis and consumption patwhays. Moreover, these studies were mostly limited to heterotrophic growth of well-studied microbes, using either glycolysis or gluconeogenesis. In this paper, we develop an unbiased computational method for the identification of the precursor set and basic bioenergetic parameters from GSMMs of microbes relying on different catabolic modes.

This paper is organised as follows. After introducing the associated concepts of our method, we apply it GSMMs. We optimise such models with FBA for maximal energetic efficiency, i.e. maximal yield of biomass on the energy source. We decompose the resulting optimal flux distribution into catabolism and anabolism and determine the identity and fluxes of the exchanged charged energy equivalents. Next, we decompose catabolism into energy and precursor catabolism to understand their contributions to the synthesis of charged energy equivalents and identify the set of used precursor metabolites. Finally, we assess the fraction of charged energy equivalents made in energy and precursor metabolism, which precursor synthesises consume charged energy equivalents, the total ATP requirement per mol of biomass, and the variation of those quantities across conditions and species varying in their mode of growth (heterotrophic, autotrophic, lithotrophic, etc.). We find variation of *Y*_*X/ATP*_ between 4.1 and 13.7 gram dry weight per mol ATP, the fraction of ATP produced by precursor and energy catabolism varies from 0 and 1, and precursor sets vary greatly between species and conditions (they only all share acetyl-CoA).

We conclude that heterotrophic growth is the most complicated and diverse case. Catabolic synthesis of charged energy-equivalents and precursor metabolites is then intertwined and dependent on conditions. We conclude that this happens because maximization of the yield of biomass on the carbon (energy) source implies minimization of catabolic-product formation and, therefore, maximization of the yield of energy equivalents during precursor biosynthesis. The optimal states we study are the optimal compromises between these objectives.

## 2 Results

### 2.1 Decomposing metabolism into energy catabolism, precursor catabolism and anabolism

Our method starts from a GSMM of a microbe and an energy source. We vary the energy source in this paper and keep the other sources of chemical elements fixed (Table S2). The metabolic state with maximal energy efficiency is calculated using FBA, by fixing the growth rate and minimising the uptake flux of the energy source. We therefore maximize the yield of biomass on the energy source (*Y*_*X/ES*_). We do not consider any minimal or maximal flux bounds, only flux-irreversibility constraints. A maximal yield solution, obtained in this manner with a single flux constraint in FBA, ensures that the optimal solution flux vector is an elementary flux mode [25]. In our method, the state of maximal energetic efficiency coincides with the state of maximal biomass yield on the energy source (*Y*_*X/E*_). This implies that growth rate is limited by nutrient availability rather than by an intracellular capacity constraint. This optimal state corresponds, for instance, to the metabolism displayed by *Escherichia coli* (*E. coli*) and *Saccharomyces cerevisiae* (*S. cerevisiae*) in an aerobic, glucose-limited chemostat when they fully respire glucose (the energy source) into carbon dioxide and water and do not make any overflow products; since, making an overflow product would always reduce the biomass yield on the carbon source.

In summary, in the case of heterotrophic growth, we aim to find the maximal value of *Y*_*X/CS*_ (with CS=ES). This involves finding the optimal compromise between producing the least amount of catabolic product, achieving maximal yield of energy equivalents on the energy source, and achieving maximal yield of precursor metabolites on the energy source. Since catabolic product formation is minimized by maximizing the yield of energy equivalents during precursor biosynthesis, the synthesis of charged energy equivalent and precursor metabolites will be intertwined. We aim to figure out how this depends on conditions.

To achieve this, we decompose the optimal flux vector into a catabolism (CAT) and anabolism (ANA) flux vector. (This method is explained in the Appendix A) Our method is upto here the same in spirit as that of Mori et al. [26]. Whereas they however had to assign a reaction as exclusively catabolic for their flux decomposition to work, our method does not require this. This allows us to focus more easily on non-model microbes. Our and Mori et al.’s method lead to the same catabolic and anabolic decomposition with respect to ATP (without considering precursors and other energy equivalents). We subsequently decompose catabolism further (than Mori et al.) into energy and precursor catabolism to highlight the importance of the latter for rationalising charged energy equivalents (i.e. ATP) yields and anabolic cost differences between microbial metabolisms.

The main function of catabolism (the fueling pathway) is to charge energy equivalents (i.e. make ATP, NAD(P)H) to ‘fuel’ anabolism. The function of anabolism (i.e. biosynthesis, polymerisation and assemblage) is to make building blocks (e.g. amino acids, nucleic acids, and lipids) and polymerise them into macromolecules (D/RNA, protein, extracellular polysaccharides and storage compounds). Anabolism consumes the charged energy carriers and returns them uncharged to catabolism for recharging. When the energy source is also the carbon source, catabolism also synthesizes precursor metabolites from the carbon source that are consumed by anabolism for the synthesis of building blocks.

Accordingly, we define catabolism as the set of reactions and reactants that together convert the energy source into charged energy equivalents and catabolic product or, when the energy source is also the carbon source, in addition into precursor metabolites. Accordingly, the precursor metabolites are a subset of the catabolic reactants. For instance, when photons (during photosynthesis) act as the energy sources, catabolism does not supply precursors, only charged energy carriers. In such cases, which we also consider below, precursor biosynthesis is anabolic and the carbon source is consumed exclusively by anabolism and precursor catabolism does not exist.

An example of a metabolism decomposition into catabolism and anabolism is shown in Figure 1. It shows the state of maximal energetic efficiency for anaerobic growth of *E. coli* with glucose as sole carbon and energy source. Catabolism consists of glycolysis and the formation pathways of three catabolic products: acetate, ethanol and formate. Seven precursor metabolites are produced by catabolism, i.e., F6P, DHAP, G3P, 3PG, PEP, PYR and ACCOA. NADH and ATP are supplied by catabolism to anabolism. The uncharged energy equivalents (NAD^+^ and ADP) and CoA are returned to catabolism by anabolism. We note that these 7 precursor metabolites are not the suggested 12 precursors by Neidhardt et al.[1].

**Figure 1:**
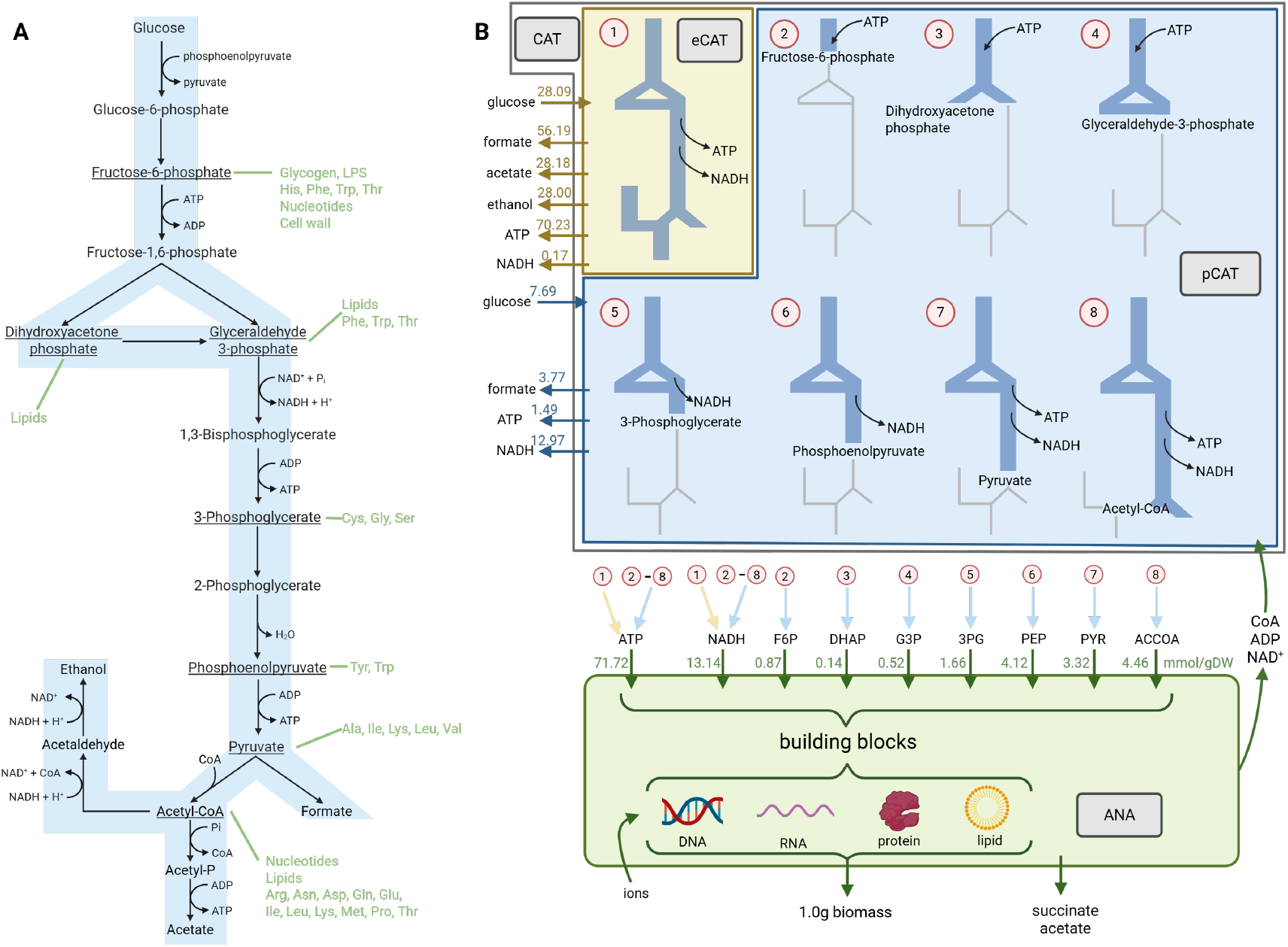
Energy catabolism, precursor catabolism and anabolism decomposition of the metabolic state of maximal energetic efficiency of Escherichia coli (E. coli), growing anaerobically with glucose as sole carbon and energy source. We computed the state of maximal energetic efficiency by setting the growth rate to 1 hr^−1^ and minimizing the glucose uptake rate (minimum = 35.78 mmol glucose gramDW^−1^ hr^−1^). After the application of our decomposition method we obtain: **A**. the catabolic network supplying charged energy equivalents and precursors for anabolism and producing the catabolic products formate, acetate and ethanol. These are made in a 1:1:2 stoichiometry when only ATP (and no NADH) is supplied to anabolism. Building block molecules that can be made from the precursor molecules are shown in green. **B**. The energy catabolism (eCAT), precursor catabolism (pCAT) and anabolism (ANA), the identity of the exchanged intermediates and flux values. The exchanged energy carriers are ATP and NADH. Anabolism returns the uncharged carriers to catabolism, i.e. CoA, ADP, and NAD. The numbers above the arrows indicate the flux values in mmol gram^−1^ hr^−1^. Energy catabolism consumes 78% of the glucose, makes 86% of the ATP and 1.0% of the NADH and the rest is done via precursor catabolism. In Section B.2, all macrochemical equations for the dissection of *E. coli* growing aerobically on glucose are given. Abbreviations: F6P = fructose-6-phosphate, DHAP = dihydroxyacetone-phosphate, G3P = glyceraldehyde-3-phosphate, 3PG = 3-phosphoglycerate, PEP = phosphoenolpyruvate, PYR = pyruvate, ACCOA = acetyl-CoA.

We subsequently split catabolism into energy catabolism (eCAT; yellow) and precursor catabolism (pCAT; blue) (Figure 1B). In eCAT, the energy source is used to charge energy carriers, and since the energy source is now also the carbon source, the carbon source is completely converted into the catabolic products. eCAT therefore consists always out of all the reactions and reactants of catabolism. None of the precursors are exchanged with anabolism by eCAT. This is done by pCAT. pCAT consists in this case out of 7 subnetworks, each a subnetwork of eCAT and producing one of the seven precursors.

Figure 1B shows a summary of the detailed exchange between between catabolism and anabolism. About 85% of the anabolic NADH demand is supplied by pCAT and the remainder by eCAT or in ANA. In contrast, 86% of the ATP is supplied by eCAT and about 80% of the consumed glucose is converted into catabolic products for the production of ATP and NADH. Three of the seven precursor metabolites require an ATP investment (F6P, DHAP, G3P), two are ATP neutral (3PG, PEP) and 2 are ATP producing (PYR, ACCOA). This leads to a net production of ATP in precursor biosynthesis. The net behavior of catabolism therefore depends on the integration of precursor synthesis and energy conservation.

In Figure 1B, anabolism (ANA; green) is also shown. It converts the 7 precursors and charged energy equivalents made by catabolism into biomass components. The majority of the charged energy equivalents is produced by eCAT. About 20% of the produced ATP is synthesised in pCAT. Part of the consumed glucose (Figure 1B) is converted completely into the catabolic products and used solely for energy conversation (catabolic glucose) whereas the remainder is used for precursor biosynthesis (precursor-related glucose).

The sum of the flux distributions of eCAT, pCAT and ANA equals the optimal flux distribution calculated with flux balance analysis (see Appendix).

### 2.2 Relations between ATP yield and requirement parameters

Our next objective is to coarse grain the catabolism (CAT, eCAT and pCAT) and anabolism pathways into net stoichiometric models, using macrochemical equations, to be able to calculate *Y*_*X/AT P*_ 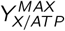 and arrive at equations used in phenomenological models of microbial ecology [24, 8] and biotechnology [27, 7]. In this section, we focus on ATP to keep the exposition simple. The computational method considers the exchange of all other possible energy equivalents (incl. ATP, NADH, NADPH and ferredoxin) and computes the ones used.

We start with the macrochemical equations of catabolism and anabolism. In biotechnology and microbial ecology, they are estimated with thermodynamics, stoichiometry and heuristics based methods [24]. We determine them here in an exact manner from a flux vector (calculated with FBA, see section A.4). This can be done for the whole cell, catabolic and anabolic flux vectors and gives the macrochemical equations of these process. Examples of such equations can be found in section B.2 of the Appendix.

The macrochemical equations of catabolism and anabolism relate the ATP formed by eCAT to the anabolic ATP demand (see Box 8 in Kleerebezem & Van Loosdrecht [24]),

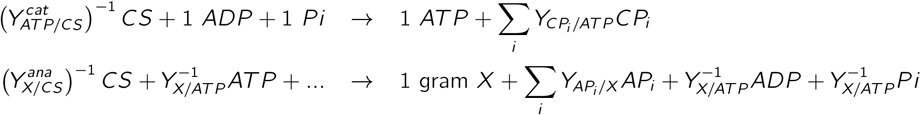

The dots denote the remaining sources of chemical elements required for biosynthesis of biomass, such as sulphate, ammonium or phosphate. *CP* and *AP* respectively denote catabolic and anabolic products. Note that the anabolic equation in this case describes the sum of pCAT (synthesis of precursors from the carbon source) and ANA (synthesis of biomass from precursors). The stoichiometric coefficient 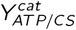 equals the catabolic yield of ATP on the carbon source. We often know its value from basic stoichiometric knowledge of a catabolic pathway. For instance, for the example shown in Figure 1A, 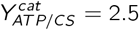, since 2.5 mol of ATP is made from 1 mole of glucose when it is completely converted into the catabolic products.

The macrochemical equation of growth is obtained by multiplying the catabolic reaction with (*Y*_*X/ATP*_)^−1^ and the addition of the resulting equation to the anabolic reaction,

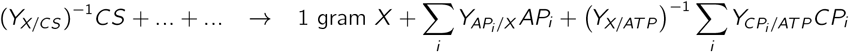

This equation equals the equation originally obtained from the FBA simulation for optimal biomass yield on energy source. The yield of biomass on the carbon source *Y*_*X/CS*_ that appears in this equation equals

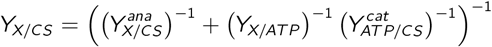

This last equation indicates that the yield increases when more ATP is made per unit carbon source and if biomass synthesis demands less ATP.

These equations indicate the central role of *Y*_*X/ATP*_ in estimations of biomass and catabolic-product yields in microbial ecology and biotechnology [24, 7, 11]. Estimating the value of *Y*_*X/ATP*_– or its constancy across species – is therefore a common objective.

To address the variation of *Y*_*X/ATP*_ and introduce the precursor metabolites, we define the ATP balance (Figure 2A) and identify their contributions. The terms in this balance are outcomes of our computational method. It corresponds to the following equation and defines *Y*_*X/ATP*_,

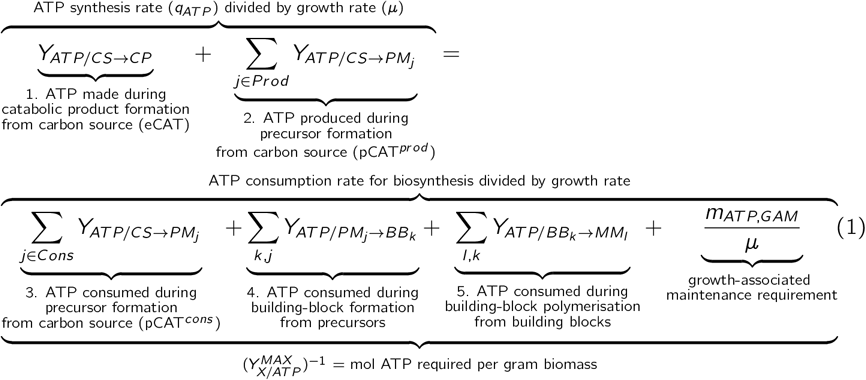

**Figure 2:**
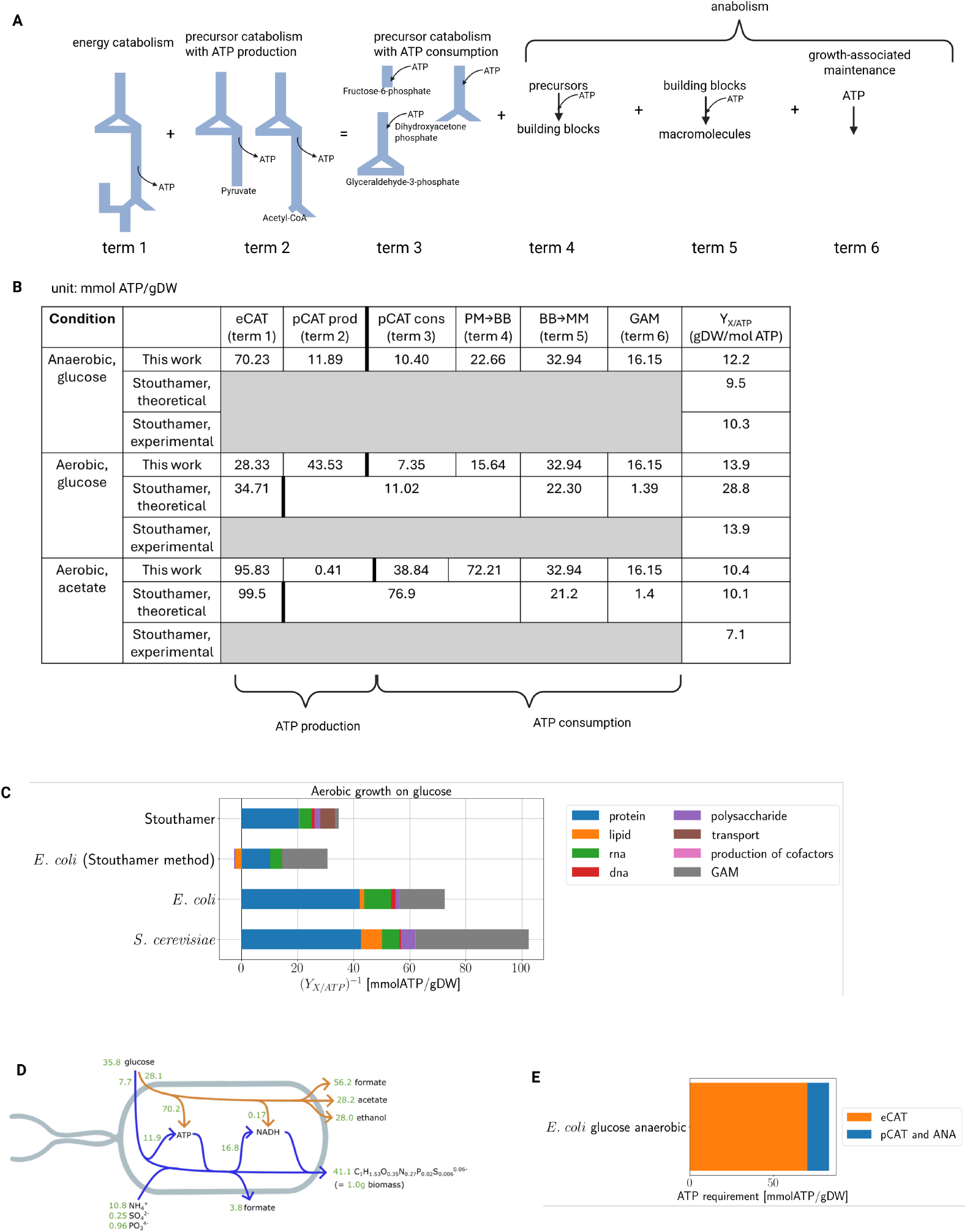
Total ATP requirement per 1 gram biomass and ATP fraction produced by energy and precursor metabolism. **A.** Schematic depiction of the ATP production and consumption balance of a cell according to equation 1. **B**. Table showing the mmol ATP supplies and demands per gram biomass for aerobic and anaerobic growth by *E. coli* on glucose. **C**. Barplot showing the ATP costs in mmol ATP/gram for aerobic growth on glucose for *E. coli* and S. cerevisiae. (The numbers in this bar graph correspond to the inverse of Stouthamer’s values multiplied by 1000.) The first bar corresponds to the calculation of Stouthamer, the second bar to our methods where we make Stouthamer’s ‘mistake’, the third bar corresponds to our method giving rise to the experimentally measured value, and the fourth bar the estimate for *S. cerevisiae*, also a model microbe, under the same conditions. **D**. A schematic depiction of the charged energy equivalent exchanges between catabolism and anabolism for anaerobic growth of E. coli and glucose. **E**. A bar plot showing total ATP production by the cell and the contributions by eCAT and pCAT.

This ATP balance was obtained by dividing the steady-state flux balance of ATP by the corresponding cellular growth rate. Term 1-3 together make up the fueling pathway and term 4-6 represent respectively biosynthesis, polymerisation and assemblage. Term 6 corresponds to the growth-associated ATP maintenance flux divided by the growth rate.

We note that we will sometimes refer to term 4-6 as anabolism and that this is a different definition of anabolism that was used in the macrochemical equations above; there it corresponded to term 2-6.

The second and third term refer to the precursor synthesis pathways that respectively produce or consume ATP. An explicit example of these contributions is shown in Figure 1B. There is net production of ATP during the synthesis of the precursors PYR and ACCOA, and net consumption of ATP during the synthesis of F6P, DHAP and G3P.

The fifth term in this equation can be estimated with the greatest accuracy and is known from basic molecular biology [9, 2, 4]; from the macromolecular composition of 1 gram biomass, the ATP equivalent costs of the chemical polymerisation bonds (in D/RNA, protein, storage compounds etc.), and the number of such bonds to be formed to make the cell’s macromolecules. Since the second to fourth term are dependent on the identity of the precursors, they are determined by the expressed metabolism. Those terms are therefore dependent on the precise conditions and the metabolic behavior of the species. When Neidhardt et al., Stouthamer and Umbarger [2, 4, 1] made their estimates of the nutrients and ATP costs to make 1 gram biomass, they assumed particular precursors and associated ATP requirements and gains. They therefore estimated the second to fourth term as a lumped sum in this equation.

The first term is not a straightforward estimate. It requires estimation of how much ATP is to be formed by catabolic product formation and results from the remainder of the ATP requirement after the second term (precursor production) has produced some of the ATP to balance with the right hand side of this equation. The first term can therefore not be estimated straightforwardly in any envelope calculation without knowledge of the fluxes through pCAT’s subnetworks, each making a precursor.

The second to fourth term in this equation vary with the precursor set (the fueling pathway and its energetic yields) and conditions. The maximization of the energetic efficiency of the GSMM will determine the optimal set corresponding to the state of maximal energetic efficiency.

Genome-scale stoichiometric models do not suffer from any of these previous three limitations and, therefore, provide more unbiased and reliable estimates.

### 2.3 The ATP balance of E. coli metabolism

We calculated the ATP requirement to make 1 gram biomass ((*Y*_*X/ATP*_)^−1^) for reference conditions of *E. coli* and *S. cervisiae* in Figure 2B-E and in Figure 3 for additional microbes and varying conditions.

**Figure 3:**
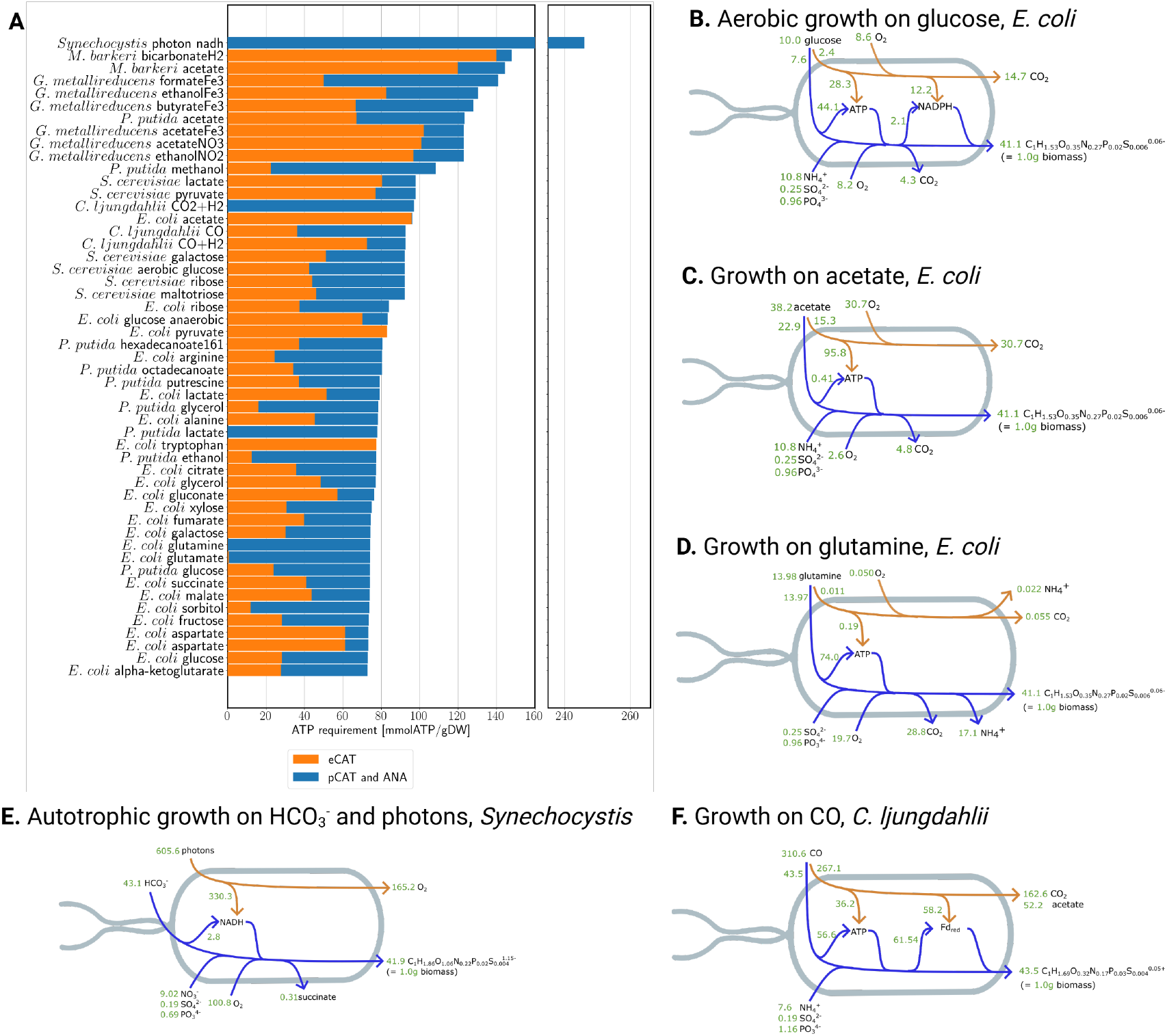
Species diversity in total ATP requirement per gram biomass and ATP-synthesis contributions of eCAT and pCAT. **A.** The mmol ATP per gram biomass requirement and the ATP synthesis contribution by eCAT and pCAT are shown across conditions. A comprehensive list of the performed simulations is given in Table S2. **B**. Aerobic growth on glucose by *E. coli* below the critical growth rate for overflow metabolism. **C**. Aerobic growth on acetate by *E. coli*. **D**. Aerobic growth on glutamine by *E. coli*. **E**. Growth of the cyanobacterium *Synechocystis* on photons as energy source and carbon dioxide as carbon source. **F**. Growth *Clostridium ljungdahlii* on carbon monooxide as carbon source.

Figure 2C shows the total ATP requirement per gram biomass expressed in terms of the individual macromolecules. These costs correspond to term 3-6 in equation 1. Our calculation predicts that *E. coli*, operating at a state of maximal energetic efficiency, requires about 82 and 73 mmol ATP to synthesise 1 gram of biomass under respectively anaerobic and aerobic conditions, when using glucose as a carbon and energy source. This leads respectively to 12.2 and 13.9 gram biomass made per mol of ATP and should be compared respectively to measured values of 10.3 and 13.9 gram per mol (reported by Stouthamer [3]). Our computational method indicates that the difference between those growth conditions is in in the adaptation of catabolism and the part of anabolism making building blocks from precursors (terms 1-4 of equation 1 and of Figure 2A). The ATP requirement for the conversion of building blocks into macromolecules and for growth-associated maintenance (GAM) are constant (fixed into the biomass reaction of the GSMM).

It is remarkable to see that a genome-scale model with a single flux constraint (growth rate) and a minimised glucose uptake rate is so close to experimental data (assuming that the GAM estimate is realistic). This indicates that *E. coli* may indeed operate close to a state of maximal energetic efficiency.

The values reported by Stouthamer in his 1973 paper are also shown in Figure 2B. His anaerobic estimate is remarkably close to the experimental data, but his aerobic value is a factor 2 too high. This discrepancy was subsequently referred to as the ‘Y_ATP_ problem’ and has remained unresolved ever since [15, 16]. Our estimate does not have this twofold discrepancy.

The ‘Y_ATP_ problem’ has to do with the treatment of the ATP gains and costs of precursor metabolism. Consider the table shown in Figure 2B. While we count all the individual contributions (term 1 to 6), Stouthamer counted production and consumption in precursor catabolism and formation of building blocks (term 2 to 4) as one term. Thus, his 11.02 mmol ATP/gram biomass term equals production in precursor catabolism - consumption in precursor catabolism - consumption for building block synthesis (term 2 - term 3 - term 4) given rise to −11.02 mmol ATP/gram, so a cost. He then determines the yield in gram biomass per mol ATP from 1*/*(34.71 *×* 10^−3^) = 28.8. (He should added the ATP costs of term 2 to do the denominator of this ratio.) He therefore underestimated the amount of ATP needed per 1 gram biomass and overestimates the yield. The following example example makes this more clear:

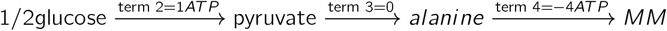

we would add a 1 to term 2, a 0 to term 3 and 4 to term 4, Stouthamer added only a 3 to the cost term, so not considering the synthesis of the 1 ATP and hereby underestimating the amount of ATP required to make 1 gram biomass.

When we perform the same calculation as Stouthamer, but using a GSMM, we obtain a similar value (31.1 gram/mol) that is also approximately twofold of (compare bar 1 to 3 in Figure 2C). Stouthamer’s error is not apparent in this calculation of anaerobic glucose and aerobic acetate, because then the term 2 values that he implicitly neglects are small relative to the term 1 values, and his use of the term 1 as the estimate of the total ATP requirement per gram biomass suffices.

Two other ways to depict the catabolic and anabolic ATP balance are shown in Figure 2D and E; both for anaerobic growth on glucose by *E. coli*. Figure 2D is a summary of the macrochemical equations of energy catabolism (orange) and precursor catabolism and anabolism lumped (blue), while Figure 2E refers to contributions of eCAT and pCAT to the ATP production (term 1 and 2 in equation 1 and in Figure 2A. pCAT is now considered in its entirety in the blue arrow in Figure 2D; so the pCAT and ANA blocks of 1B are fused into 1 process that only consumes charged energy equivalents. Figure 2E shows how much ATP is produced in eCAT and in pCAT (plus ATP production by a few anabolic reactions, always negligible and less than 2.5% see Fig S6), the sum equals the total production of ATP per 1 gram biomass.

### 2.4 Catabolic diversity: energy carriers

Figure 2B indicates that during aerobic growth on glucose most ATP is made in precursor catabolism while the opposite is the case under anaerobic conditions (table in Figure 2). Since the reactions composing catabolism depend on the growth conditions, we tested whether the total ATP requirement and the fraction produced by precursor metabolism varies between microbial species and conditions.

We calculated the eCAT and pCAT contributions to total ATP synthesis across 50 growth conditions and 7 microbial species (Figure 3A). The relative amount of ATP synthesised in eCAT versus pCAT, as well as the total amount of ATP synthesised vary greatly from 72.8 - 246.1 mmol ATP per gram biomass (Figure 3B). In some cases, almost the entire anabolic ATP requirement is produced during precursor synthesis (*E. coli* glutamine); while in other cases, no ATP is produced during precursor synthesis at all (*E. coli* pyruvate, acetate).

To understand this better, Figure 3B-F shows detailed examples of several cases shown in Figure 3A. The fraction of ATP produced during precursor biosynthesis on aerobic growth by *E. coli* on glucose and growth on acetate are 0.60 (Figure 3B) and 0.004 (Figure 3C), respectively. These differences can be explained by considering the catabolic network. Consider the production of pyruvate when growing on glucose: this is accompanied by the production of 1 pyruvate, 1 ATP, and 1 NADH. In oxidative phosphorylation, this NADH can be oxidized for the formation of 1.75 ATP. Thus, in total 2.75 ATP/pyruvate are formed that do not need to be produced by fully converting glucose into CO_2_. During growth on acetate, energy is conserved in the TCA cycle. No carbon precursors are then formed, since energy generation is accompanied by the release of two molecules of CO_2_ (all the carbon atoms in acetate). The glyoxylate shunt and gluconeogenesis convert acetate, which does not involve a net ATP synthesis, but, instead an ATP cost; as acetate is first converted into acetyl-CoA, at the expense of an ATP.

In Figure 3D, *E. coli* uses glutamine as its sole carbon, nitrogen and energy source. In this case, the majority of its ATP production runs via precursor metabolism. The formed catabolic products are ammonium and carbon dioxide, and eCAT’s contribution to glutamine uptake is negligibly small. In the final section, we return to this case when we discuss the precursor metabolite sets and the associated ATP production.

Cyanobacterium *Synechocystis* sp. PCC 6803 growing autotrophically, using photosynthesis with photons as its source of energy is shown in Figure 3E. All ATP synthesis is synthesized by anabolism, because the exchanged charged energy carrier is NADH. NADH is a more sensible choice in this case. If ATP would have acted as the exchanged energy carrier, the catabolic macrochemical equation consists only of consumption of photons with production of ATP (Eq. 2), i.e.,

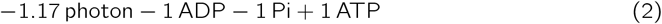

With NADH as exchanged energy carrier, H_2_O is consumed and O_2_ is produced in catabolism (Eq. 3) leading to

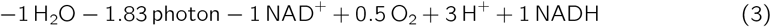

The latter process is a much better reflection of the underlying mechanistic biochemistry. In the light-dependent reactions, photons excite the electrons donated by water. These excited electrons subsequently drive proton movement across the membrane in the electron transport chain. These exported protons are subsequently used by ATP synthase to form ATP. Finally, the electrons are donated to NAD^+^ to form NADH [28]. Thus, no ATP can be generated without the electrons from water. These two processes are however not stoichiometrically coupled in the genome-scale model of this microbe. To synthesise biomass, both ATP and NADH are required. However, because the two are not linked in the genome scale model, there is no way to determine how much ATP versus NADH is required. Therefore, NADH as the energy carrier is the most sensible choice, which we enforced in this case. This case illustrates the importance of the choice of energy carrier. This issue is discussed further in Section B.4.

The anaerobic acetogen *Clostridium ljungdahlii* (*C. ljungdahlii*) can grow on combinations of CO, CO_2_ and H_2_ as energy donor. This microbe can use either CO_2_ and H_2_ as substrates, or CO, with or without H_2_ (Figure 3F). It relies on the Wood-Ljungdahl pathway, which converts 2 molecules of CO into acetyl-CoA leading to either acetate and CO_2_ (or acetate and ethanol) as catabolic products [29]. The Wood-Ljungdahl pathway harvests some energy, but additional H_2_ can donate electrons to energy carriers. When growing on only CO, we find that both ATP and reduced ferredoxin (Fd_red_) were produced by catabolism according to the following stoichiometries (Figure 3F),

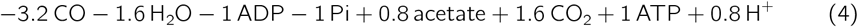

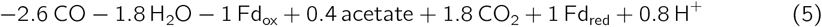

When both CO and H_2_ are used as energy substrates, ethanol is produced instead of acetate, with maximal energetic efficiency of CO is the FBA objective. In this case, either Fd or ATP be defined as energy carrier, Eq. 6 and Eq. 7, respectively. When Fd is defined as an energy carrier, H_2_ is produced as a by-product in anabolism. As the overall process consumes H_2_, we therefore chose ATP as exchanged energy carrier.

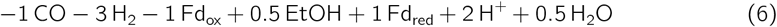

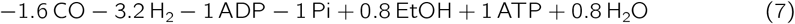

Lastly, when *C. ljungdahlii* grows on CO_2_ and H_2_, Fd must be defined as energy carrier (according to Eq. 8), and ATP cannot be defined as energy carrier, because the balanced production of acetate from CO_2_ and H_2_ does not have net ATP production. As a consequence, no ATP production occurs in energy catabolism (Figure 3A). ATP is produced anabolically via a proton gradient using ATPase.

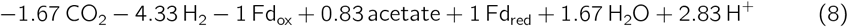

Since ATP is in this latter case not exchanged, the fraction of ATP that is produced in anabolism (and again consumed in anabolism) is clearly 1, as there must be balance within each subsystem. For growth on CO alone, the anabolic ATP fraction was 0.61, and for growth on CO and H_2_, the fraction was 0.22.

These three examples show that not only the quantity of energy carrier that must be produced by anabolism can be variable, but also which energy carrier is used, even in relatively similar growth conditions.

This section indicated that the role of energy catabolism, both qualitatively (which energy carriers are produced) and quantitatively (the production fractions of the energy carriers), shows significant variability between species and conditions. This variation depends on details of the biochemistry described in the associated genome-scale stoichiometric models. In the next section, we consider the effect of precursor metabolism on this diversity.

### 2.5 Integration of charged energy carriers (identity and demand) and precursor metabolism for E. coli

The previous sections indicate that the identity of charged energy carriers and their relative production by eCAT and pCAT varies between conditions. In Figure 4, we provide a more comprehensive overview of *E. coli* at states of maximal energetic efficiency on 23 different carbon sources.

**Figure 4:**
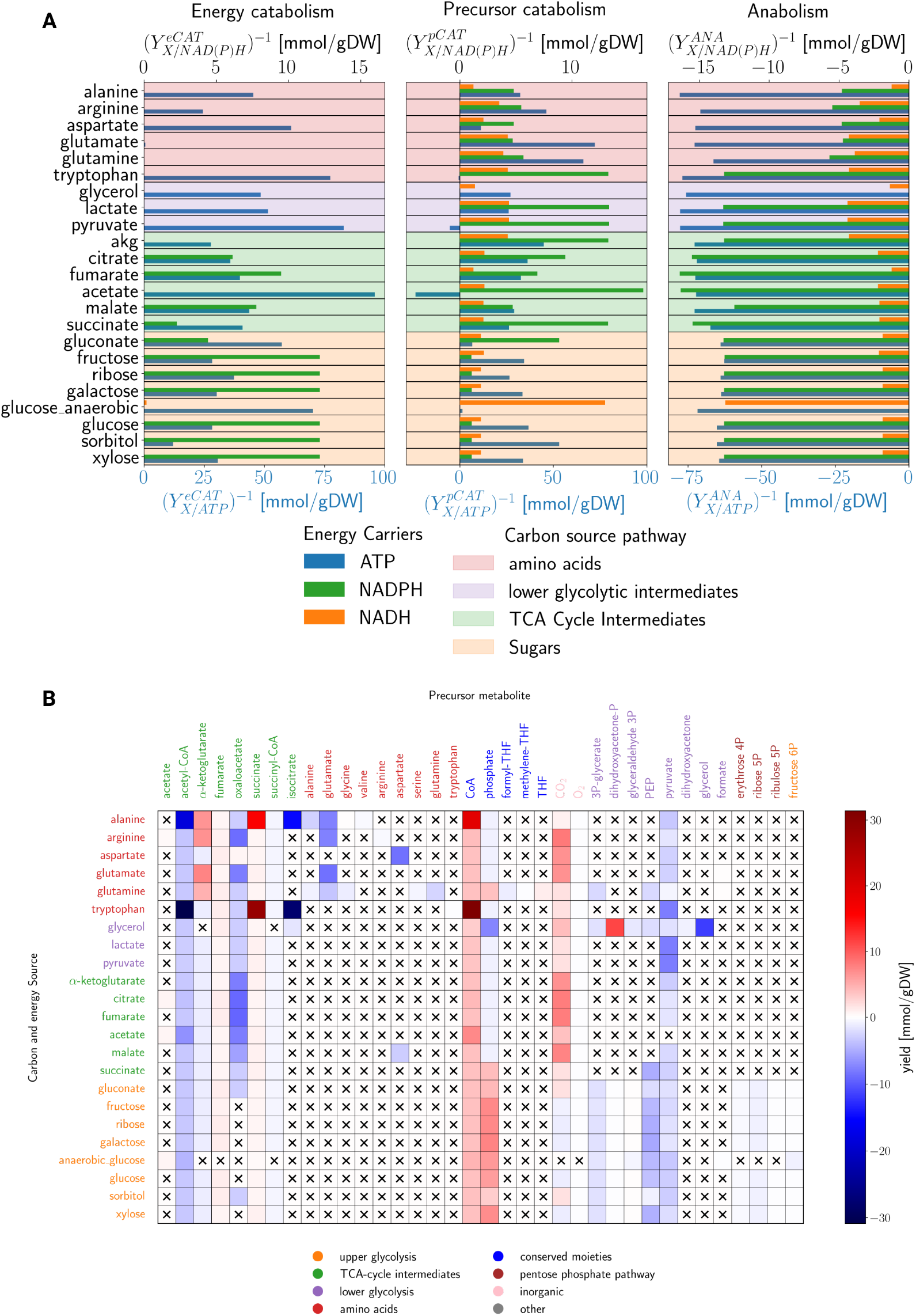
E. coli’s variation in exchange and production of charged energy carriers and precursor usage. **A.** The total amount of ATP, NADH and NADPH produced or consumed in energy catabolism, precursor catabolism and anabolism across 23 different energy sources of *E. coli*. **B**. Overview of the consumed precursor metabolites and anabolic products produced by anabolism across the 23 different energy source considered in Figure A. The color indicates the stoichometric coefficient in the macrochemical equation of anabolism. An overview of the perforemd simulations in *E. coli* is given in Table S2.

In Figure 4A, the production or consumption of NADH (green, top axis), NADPH (orange, top axis) and ATP (blue, bottom axis) is shown by eCAT (left figure), pCAT (middle figure) and ANA (right figure). The sum of each bar across the three figures equals zero, indicating balanced production by CAT and consumption by ANA.

Energy catabolism exchanges ATP and NADPH with anabolism when *E. coli* grows on sugars and TCA cycle intermediates, otherwise only NADH. Only during anaerobic growth on glucose does it exchange NADH and ATP. Precursor catabolism is a much more diverse supplier than energy catabolism, supplying always all three charged energy carriers. It consumes ATP during growth on pyruvate and acetate. Anabolism has the largest demand for ATP across all the conditions, which from 62.7 - 77.8 mmol ATP per gram biomass.

Sugars have the lowest anabolic energy requirements. The differences stem from the differences in precursors. Considering production of DNA from glucose, anabolism has to convert fructose-6P into DNA, while, when growing on acetate, anabolism has to convert oxaloacetate into DNA, via the ATP-consuming gluconeogenesis. The fraction of ATP that is produced during biomass production is at most 0.025 (Figure S6), and the ATP requirement of ANA is fairly constant (Figure 4). Thus, the variability in the quantity of energy harvesting stems from the precursor production.

The number of precursors used across the conditions varies significantly between conditions (min: 6, mean: 9.6, median: 8, max: 14). For substrates with a larger catabolic pathway, e.g. glucose (glycolysis, pentose phosphate pathway and TCA cycle), the number of precursors was larger than for substrates with a shorter catabolic pathway, e.g. acetate (only the TCA cycle). From the macrochemical equations of anabolism for different carbon substrates, starting from the precursors, the identity and requirement of the precursors were determined (Figure 4B).

Depending on the place where the energy source enters central carbon metabolism, some observations can be made. Generally, more oxaloacetate is consumed for growth when a TCA cycle intermediate is the substrate, as oxaloacetate is the metabolite that is consumed for gluconeogenesis. In addition, PPP intermediates are only carbon precursors when growing on sugars, as the PPP is then a catabolic pathway that is related to the production of NADPH. When an amino acid is the energy source, such that amino acids are generally also a carbon precursor metabolite - they are not have to be synthesized from another pure carbon precursor.

Across conditions, we also find similarities in the precursor sets: acetyl-CoA and pyruvate are (almost) universal precursors. Fumarate and succinate are always byproducts from anabolism, produced using the fumarate/succinate redox couple and as a product of succinyl-CoA consumption, respectively.

### 2.6 Catabolic diversity: precursor metabolism

In Figure 5, we show simplified metabolic maps of the precursor catabolisms of different species and conditions. We will highlight a few interesting cases and differences. Because *E. coli* relies on (canonical) EMP-glycolysis and *Pseudomonas putida* (*P. putida*) relies on Entner-Doudoroff glycolysis, which partly uses the pentose phosphate pathway, their precursor metabolites differ. Both organisms use the same TCA cycle intermediates as precursors. When these microbes grow on acetate the precursor metabolites are very similar and the differences between the exchange charged energy equivalents are minor (*E. coli* exchanges more of them but at relatively negligible values). Growth on this same energy source (acetate) by *Methanosarcina barkeri* (*M. barkeri*) is, however, very different from *P. putida* and *E. coli*. The only common precursor is acetyl-CoA. The environments are different now: *P. putida* consumes O_2_ while *M. barkeri* produces CH_4_ anaerobically. They perform the energy-harvesting conversions using completely different pathways, which is why they do not share many precursors. In another strictly anaerobic environment, *C. ljungdahlii* can consume CO to form CO_2_ and acetate. The precursors that are used in this situation are again very different from the ones used by *M. barkeri* (Figure 5D). Again, they do both use acetyl-CoA and CO_2_ as a precursor. This shows that the precursors consumed depend on the catabolism of the organism.

**Figure 5:**
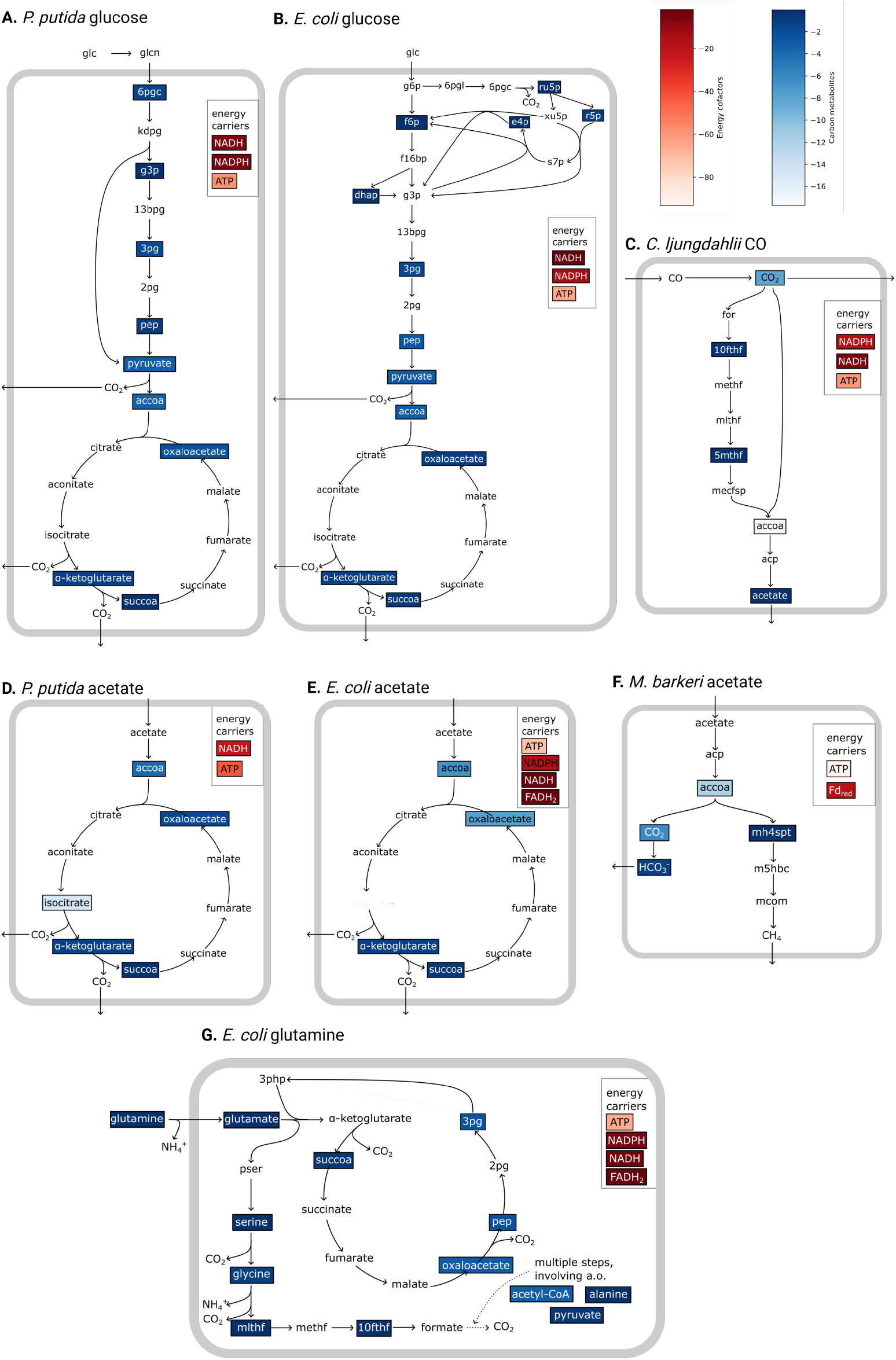
Diversity of selected precursor metabolisms across species and conditions. The precursor metabolites are shown in blue and the exchanged energy equivalents are shown in red the box. Their red and blue levels indicate relative usage by anabolism. **A**. *Pseudomonas putida* growing aerobically on glucose,**B**. *Escherichia coli* growing aerobically on glucose, **C**. *Clostridium Ljungdali* growing aerobically on glucose, **D. E**. *Pseudomonas putida* growing aerobically on acetate, *Escherichia coli* growing aerobically on CO, **F**. *Methanosarcina barkeri* growing aerobically on acetate, **G**. *Escherichia coli* growing aerobically on glutamine. Abbreviations of metabolites are BiGG identifiers [30].

Thus, the carbon precursors required for growth are highly diverse and depend on the metabolic pathways of the species. This is a consequence of precursor metabolites being the reactants of the energy catabolism. If the energy catabolism pathways of two microbes share only a few metabolites then it is likely that their precursor metabolites vary as well.

Neidhardt et al. [1] took another perspective and argued that a fixed set of 12 precursor metabolites are always used: glucose-6-phosphate, fructose-6-phosphate, ribose-5-phosphate, erythrose-4-phosphate, triose-phosphate, 3-phosphoglycerate, phosphoenolpyruvate, pyruvate, acetyl-CoA, *α*-ketoglutarate, succinyl-CoA, and oxaloacetate. (Stouthamer [2] and Umbarger [4] used very similar precursor sets.) These are metabolites that occur in glycolysis, the pentose phosphate pathway and the TCA cycle. It is likely that many non-model microbes may lack some of these intermediates, because they exploit different metabolic pathways. But even for *E. coli* it is doubtful that this list is constant across conditions. We did not recover it completely in any single case. When we enforced growth on these 12 precursors while the catabolic pathway (defined for aerobic growth on glucose) was deactivated, growth was infeasible in the active network (for details see Appendix B.8).

## 3 Discussion

This work indicates that the diversity of heterotrophic microbial metabolism – the most complicated case – is in the reactions that compose fueling pathways (catabolism) and biosynthetic pathways (converting precursors into building blocks). The molar ATP requirement (and of charged energy equivalents in general) per gram of biomass (in eq. 1) consists of a constant requirement (polymerisation and assemblage), and a variable component, associated with energy conservation and precursor synthesis from the carbon source and their conversion into building blocks (Figure 2B). Since these processes are species and condition dependent, the parameters *Y*_*X/AT P*_ nor 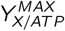 (eq. 1), are indeed not constants [3]. We decomposed them into separate distributions to make this more clear. While doing so, we solved the *Y*_*ATP*_ problem associated with Stouthamer’s calculations.

Most of the non-intuitive results of this work derive from the dual function of catabolism in the case of heterotrophic growth, i.e. the associated set of precursor metabolites and its contribution to consumption of synthesis of charged energy equivalents. We believe that this set varies, and therewith the biosynthetic pathways, because we focus on the metabolic state of maximal energetic efficiency, corresponding to the maximal yield of biomass on the carbon (energy) source. This state is the state of carbon-limited growth in the chemostat. This optimal state is the best compromise between the following “forces”. Since catabolic product formation (eCAT) yields only charged energy equivalents, and instead of precursor metabolites, a catabolic product, it constitutes a waste of the carbon source and should be minimized. This implies that the charged energy equivalent synthesis during precursor synthesis from the carbon source (by pCAT) is maximized, and that therefore the precursor biosynthesis pathways are chosen that yield the most ATP. Finally, the charged energy equivalent consumption by the biosynthetic pathways, converting precursors into building blocks, is minimized to minimize catabolic product formation. Since these objectives are interdependent and biochemically intertwined, we believe that this underlies the diversity of the optimal metabolisms that we found in the paper and results from Figure 2–5.

We note that none of our computations led to the 12 precursors postulated by Neidhardt, et al. [1], not even when glucose was the carbon and energy source. We have the impression that that set is indeed a feasible precursor set but not the one that corresponds to the state of maximal energetic efficiency.

The state of maximal energetic efficiency coincides with the state of maximal growth rate when growth is limited by nutrient availability and not by an intracellular capacity constraint. These conditions can be attained in nutrient-limited chemostats below the critical growth rate where overflow metabolism commences. Our results do therefore not apply to all cases of growth. In particular not for nutrient excess conditions when respirofermentative growth occurs, as fermentative growth is generally a state of suboptimal energetic efficiency. Although predicting the charged energy equivalent balances and the precursor set can be done with the method presented in this paper, these states are harder to predict because it is still an open problem at which growth rate overflow metabolism starts and how much it contributes during optimal batch growth at nutrient excess. Although this can be predicted with proteome-constraint models [31, 32], we lack those for non-model microbes. We note that the state of maximal energetic efficiency may also apply to human metabolism but the cellular overarching objective is not growth rate but carrying out the function of the cell type of interest.

Our results depend on how we define catabolism, as this determines the set of candidate precursor molecules. In our definition, catabolism constitutes the metabolic network that completely converts the carbon source into the catabolic products and charged energy equivalents. Thus, all the chemical elements (not all electrons) of the carbon source end up in the catabolic products. The precursor metabolites are then chosen from the reactants in this reaction pathway. The catabolic pathway is an outcome of the optimization of a whole-cell physiological parameter, i.e. the biomass yield on the carbon (energy) source and can therefore only be deduced after a FBA computation.

Our formalism is based on the concept of elementary flux modes [25]. These are flux distributions of a metabolic network that have no redundant fluxes (no flux can be removed without loss of steady state) and one flux value suffices to determine all others. The flux distribution through a metabolic network that has the maximal yield of a network product on a network substrate is always an elementary flux mode. Thus, the network with maximal energetic efficiency that we compute is an elementary flux mode. Since they each have a single independent flux, the stoichiometric coefficients of the net conversion they catalyse (their macrochemical equation) are constant, which allowed us to make Figures 2-5 that are unambiguous about the energy equivalent and precursor requirements. We view this as a key strength of our approach. Elementary flux modes are, however, still a concept mostly known to a small number of theoreticians, despite their useful properties for quantitative physiology studies.

In relation to previous attempts [2, 4, 1, 9, 7], our method has several advantages: i. it ensures flux balance of all metabolites (balanced growth), ii. it does not require a priori defined set of precursor metabolites (unbiased), and iii. it allows for a precise definition of the reference metabolic state (by postulating a metabolic objective). It is therefore ideally suited for a comparative analysis of microbial metabolisms; in particular of poorly-characterised microbes that vary in their modes of growth (hetero-, auto- or lithotrophic).

What makes the earlier works of Neidhart et al., Forrester & Walker, Stouthamer, and Heijnen & Van Dijken [1, 2, 9, 3] so appealing (and what motivates us) is the search for stoichiometric principles of microbial growth-associated metabolism in terms of a small set of whole-cell physiological concepts. This ‘middle way’ approach [33], has proven very successful in the last decade in understanding shifts in metabolism and ribosomal usage as function of growth rate, from the viewpoint of optimal allocation of limited biosynthetic resources [34, 35]. A disappointing outcome of this work is that we do not find a simple relation, other than eq. 1, between the identity of the precursors, the fraction of charged energy equivalents made by precursor metabolism, and the ATP requirement for 1 gram of biomass. We did attempt to find such a dependency (e.g. Figures S4 and S5), but did not succeed and found no way of linking the considered parameters to the identity and properties of the precursor metabolites. We hope that this work motivates further research into this fundamental stoichiometric and bioenergetic principles of microbial growth.

## 4 Methods

We developed a method for the decomposition that is applicable to any FBA calculation of a stoichiometric model. We perform the decomposition on metabolic networks with optimal biomass yield on the energy source (which is often also the carbon source), as this is the best reference condition to compare different species. Because we minimize the use of energy source, we maximize the amount of ATP that is generated during precursor synthesis. This is how ATP production and precursor metabolism are linked. In addition, maximum ATP production in catabolism determines the identity of the precursors. This is why this is the correct way to define this.

To calculate a metabolic network with optimal biomass yield, we perform flux balance analysis on a genome scale metabolic model. We disregard non-growth associated maintenance (NGAM), as this parameter is condition-dependent [36] and decreases the biomass yield. The decomposition consists of two parts. First, the eCAT network is determined. Then, the anabolic network, i.e. carbon precursor consumption, is determined. These must be done in two separate steps, because the energy catabolism network determines the identity of the carbon precursors, as they are defined as intermediates of eCAT. In short, the following sequence of steps are involved (for details, see Appendix A).

1. Dissection based on energy carriers.
  a. **Minimize the energy source uptake for a given biomass formation flux**. This computation corresponds to the most energetically efficient mode of microbial growth with maximal biomass yield on substrate (*Y*_*X/E*_).
  b. **Remove inactive reactions**. We removed all zero-flux reactions from the model. This leads to the identification of the ‘active’ metabolic network, which is the network in which we determine the subnetworks.
  c. **Calculate anabolic stoichiometry**. In this decomposition, the anabolic subsystem consumes charged energy carriers made by catabolism and produces itself the precursors from the carbon source and then turns those into macromolecules. To determine the anabolic influx of charged energy carriers, we add synthesis reactions for all energy-carriers, e.g. *ADP* + *Pi → ATP* to the active model. Next, minimize the carbon source uptake flux given the reference growth rate. The outcome is the anabolic flux distribution with charged energy carriers and all chemical-element sources as substrates and as products: biomass, anabolic products and uncharged energy carriers.
  d. **Determine the minimal set of exchanged charged energy carriers**. We determine the variability in the usage of the energy carriers to define if they are essential. The minimal number of energy carriers was then selected for exchange (for details and an example, see Appendix).
  e. **Calculate catabolic stoichiometry**. The network for energy catabolism was calculated by minimizing the uptake flux of the energy source to produce the required energy carriers that were obtained in the previous step.
2. Further dissection based on precursors.
  a. **Define potential carbon precursors**. Any intermediate in the catabolic flux distribution is defined as a possible carbon precursor that can be consumed by the anabolic subnetwork.
  b. **Calculate the biomass-producing flux distribution** All catabolic fluxes are deactivated to prevent interconversion of carbon precursor in the biomass producing network. Then, consumption and production of the candidate carbon precursors and all energy carriers is allowed. A feasible, growing flux distribution is found by setting the biomass reaction to a value and optimizing that same flux.
  c. **Calculate the flux distribution for precursor production**. From the dissection based on energy carriers (step 1a-e), we know the energy harvesting flux distribution. The precursor catabolism flux distribution is now determined by subtracting the energy-catabolic flux distribution and the anabolic flux distribution from the total flux distribution.

This methodology results in subnetworks of energy catabolism, precursor catabolism and anabolism and their corresponding net reactant stoichiometries (macrochemical equations). From the macrochemical equation, we determine the elemental composition of biomass. Next, the desired bioenergetic parameters are calculated. The macrochemical equations for *E. coli* growing aerobically on glucose are shown in Appendix B.2. The exchange of energy carriers is robust against flux variability in the optimal solution (Appendix B.3). However, the exchange of precursors is not unique (See Appendix B.5). The variability mostly stems from some carbon precursors that are converted into one another via anabolic routes and not alternative pathways that produce biomass. Thus, we consider a single solution of the method.

### 4.1 Genome scale models

Genome scale models used for this work are: iHN637 [37] (*C. ljungdahlii*), iAF987 [38] (*G. metallireducens*), iMG746 [39] (*M. barkeri*), Yeast9 [40] (*S. cerevisiae*), iJN1462 [41] (*P. putida*), iML1515 [42] (*E. coli*) and iSynCJ816 [43] (*Synechocystis*). The models for *E. coli, S. cerevisiae* and *Synechocystis* required some adaptations (see Section A.5).

## 5 Acknowledgements

Dedication: We dedicate this work to the memory of Professor Dr A. H. Stouthamer (1931-2023), the predecessor of FJB at the Vrije Universiteit Amsterdam, who pioneered quantitative calculations of the ATP requirement for the synthesis of 1 gram biomass.

We thank Jack Pronk, Nico Claassens, Matteo Mori, Bas Teusink, Douwe Molenaar and Johan van Heerder for insightful discussions. MR and FJB acknowledge funding by NWO-XL grant OCENW.XL21.XL21.007 ‘Taking Control of Metabolism in Microbial Cell Factories by Applying Noncanonical Redox Cofactors’.

The figures were generated using BioRender.com.

