## Supplementary for "Entangled stoichiometric objectives shape microbial catabolism"

### A Supplementary methods

#### A.1 Decomposing a GSMM into catabolism and anabolic subnetworks

The aim is to disentangle an optimal flux distribution, i.e. in our case one having a maximal biomass flux over energy-source uptake flux (i.e. maximal biomass yield on energy source, $Y_{X/E}$; state of maximal energetic efficiency), into three flux distributions: energy catabolism, precursor catabolism and anabolism. Their weighted sum equals the flux distribution through the entire metabolic network with maximal $Y_{X/E}$.

We determine the solution with optimal $Y_{X/E}$ by performing a flux balance analysis (FBA) in which the biomass objective flux (BOF; growth rate) is set to 1.0 gDW gDW$^{-1}$ h$^{-1}$ and the consumption rate of energy source was minimized such that their maximal ratio was obtained (equivalently, we could have set the consumption rate of energy source to 1.0 and maximized the BOF). This optimization of the biomass yield leads to the calculation of an elementary flux mode (EFM); since only a single flux constraint is hit and the maximal yield solution is guaranteed to be an elementary flux mode [1]. (Since we optimize the value of one flux given only the value of one other flux, the solution of the FBA is a so-called elementary flux mode (EFM) when a simplex algorithm is used for this computation (when an interior point algorithm is used, a convex combination of EFMs can result that each generate the maximal $Y_{X/E}$ value).)

An EFM is a steady-state solution of the model that obeys all irreversibility constraints and is no superposition of (a set of) other EFMs [1]. Importantly, an EFM only has one flux degree of freedom, i.e. if the flux value of one flux is known, all the other fluxes scale with that value. It is also known that any metabolic network that achieves a maximal yield (which can be any flux ratio) is an EFM [1]. To obtain an FBA solution with only one EFM, i.e. removing all net-zero cycles, all reversible reactions were split into a forward and a reverse reaction (to ensure only positive flux values) and then we used a simplex algorithm to solve the linear program [2]. In addition, the non-growth-associated maintenance constraint was removed. Then, the uptake of the energy source (ES) is minimized.

The linear program detailing the optimization is given in Eq. S1 (note in the optimal state $v_{BOF} = 1$) (Figure S1, Step 1). (Note that there is no minimal value for the ATPM flux.) The non-growth associated maintenance flux was neglected to enable comparison of optimal energetic states of different conditions and organisms.

$$\begin{aligned} \text{minimize} \quad & v_{\text{uptake of energy source}} \\ \text{subject to} \quad & \boldsymbol{N} \cdot \boldsymbol{v} = \boldsymbol{0}, \\ & \forall i: \ 0 \leq v_i \leq 1000, \\ & v_{BOF} \geq 1 \end{aligned} \tag{S1}$$

The optimal solution of this linear program we refer to as $\boldsymbol{v}^{opt}$, which is formally defined as:

$$\boldsymbol{v}^{opt} = \arg\min_{\boldsymbol{v}} \left\{ v_{\text{uptake of energy source}} | \boldsymbol{N} \cdot \boldsymbol{v} = \boldsymbol{0}, \ \forall i: \ 0 \leq v_i \leq 1000, v_{BOF} \geq 1 \right\}. \tag{S2}$$

Even though we use a simplex algorithm, alternative equally optimal EFMs may still exist which can be tested with flux variability analysis. This is not a problem for our method, just as long as key energetic parameters that we are interested do not vary in this solution set (which we always tested and did not) (See Section B.3).

From $\boldsymbol{v}^{opt}$ we can determine the macrochemical equation of growth, i.e.

$$Y_{X/E}^{-1} \text{ C-source} + n \text{ N-source} + p \text{ P-source} + s \text{ S-source} + .... \rightarrow 1 \text{ gram biomass} + a \text{ anabolic products} \tag{S3}$$

using the method described in Section A.4. One can key parameter we have now determined, i.e. $Y_{X/E}$, which relates to other model parameters as,

$$Y_{X/E} = \frac{v_{BOF}^{opt}}{v_{\text{uptake of energy source}}^{opt}}, \tag{S4}$$

i.e. the quantity we maximised in the linear program. This is one of the bioenergetic parameters we are interested in this paper, the other two $Y_{ATP/E}$ and $Y_{X/ATP}$ require the determination of the macrochemical equations of the subnetworks (eq. S3).

To prepare for the determination of the subnetworks and their related fluxes, we remove all the reactions and metabolites from the network that were not used in the optimal solution of the FBA, resulting in the "active model" and its associated stoichiometric matrix, $\boldsymbol{N}_A$ and optimal solution $\boldsymbol{v}_A^{opt}$ (which is an element of $\boldsymbol{v}^{opt}$, the optimal solution of the linear program) (Figure S1, Step 2; $\boldsymbol{v}_A^{opt}$ constitutes all the non-zero entries of $\boldsymbol{v}_A^{opt}$). Note that this leaves the macrochemical equation of growth unaltered as we removed all rates from $\boldsymbol{v}^{opt}$ with zero fluxes; hence $\boldsymbol{m}^T \boldsymbol{N} \boldsymbol{v}^{opt} = \boldsymbol{m}_A^T \boldsymbol{N}_A \boldsymbol{v}_A^{opt}$ (explained in Section A.4).

The number of columns of the active stoichiometric matrix minus its (row) rank equals the number of active EFMs in the solution space. In this case, one EFM is active: so the number of columns in $\boldsymbol{N}_A$ minus its rank equals 1. In other words, the active model has one degree of (flux) freedom (i.e. 1 independent flux, given this value all other values can be determined). We aim to disentangle the biomass optimal flux distribution into three subnetworks, in two steps, as described in Section A.2 and A.3.

#### A.2 Step 1: Dissection based on energy carriers

We aim to disentangle catabolism and anabolism, given $\boldsymbol{v}_A^{opt}$ and the identification of the energy carriers, such as ATP and NADPH. We demand that the catabolic and the anabolic half reactions are each charge and chemical-element balanced.

Catabolism is defined as the production of charged energy carriers (e.g. ATP or NADPH) from their uncharged state (ADP or $NADP^+$) during the assimilation of the energy source (substrate) (as defined by the medium and the microbe's metabolism).

Next, the produced ATP from ADP and Pi (Eq. S5) or the produced NADPH from $NADP^+$ (Eq. S6) can be defined as energy carriers exchanged between catabolism and anabolism. We use the example of ATP and NADPH here, but later we explain how we determine the energy carriers that are exchanged. When one defines catabolism as producing ATP and NADPH for anabolism, an energy-carrier production reaction is added to the active model, one at a time. To determine the catabolic flux distribution, the lower bound of this energy-carrier reaction (with flux $v_{ec}$ is set to 1, the constraint on the BOF is removed, and $v_{upt,es}$ is minimized (Eq. S7). The energy carrier reaction with rate $v_{ec}$ is either

$$\mathrm{ADP} + \mathrm{Pi} + \mathrm{H}^+ \rightarrow \mathrm{ATP} + \mathrm{H_2O} \tag{S5}$$

or

$$\mathrm{NADP}^+ + \mathrm{H}^+ + 2\mathrm{e}^- \rightarrow \mathrm{NADPH}. \tag{S6}$$

Next we perform the linear program, once for ATP and once for NADPH as energy carrier (Figure S1, Step 3),

$$\begin{array}{ll} \text{minimize} & v_{upt,es} \\ \text{subject to} & \boldsymbol{N}_A \cdot \boldsymbol{v} = \boldsymbol{0}, \\ & 0 \leq v_i \leq 1000 \quad i = 1, \ldots, n, \\ & v_{ec} \geq 1 \end{array} \tag{S7}$$

In both cases, we find a flux vector solution that corresponds to a single EFM; because we are considering a linear program with a single flux constraint and this we check by verifying that the number columns of the active stoichiometric matrix minus its rank equals 1. Although we aimed to find the flux distribution to produce an energy carrier, it could turn out that in this flux distribution, biomass is also produced. This occurs when one energy carrier cannot be produced independently of another and the dependent energy carrier can only be consumed in anabolic fluxes. As we have defined biomass production to be anabolic, we correct for this by subtracting biomass production from the energy carrier, by subtracting the optimal, biomass producing flux distribution $\boldsymbol{v}_{\boldsymbol{A}}^{\boldsymbol{opt}}$, corrected for the amount of biomass produced in the energy-carrier producing flux distribution ($v_{BOF}^{ec}$), from the energy-carrier producing flux distribution $\boldsymbol{v}_{\boldsymbol{FBA,ec}}$. (Eq. S8). Note that $v_{ec,norm}$ is not necessarily an EFM, as it can contain negative fluxes for reactions that have a lower bound of 0.

$$\boldsymbol{v}_{\boldsymbol{ec,norm}} = \boldsymbol{v}_{\boldsymbol{FBA,ec}} - \frac{v_{BOF}^{ec}}{v_{BOF}^{opt}} \cdot \boldsymbol{v}_{\boldsymbol{A}}^{\boldsymbol{opt}} \tag{S8}$$

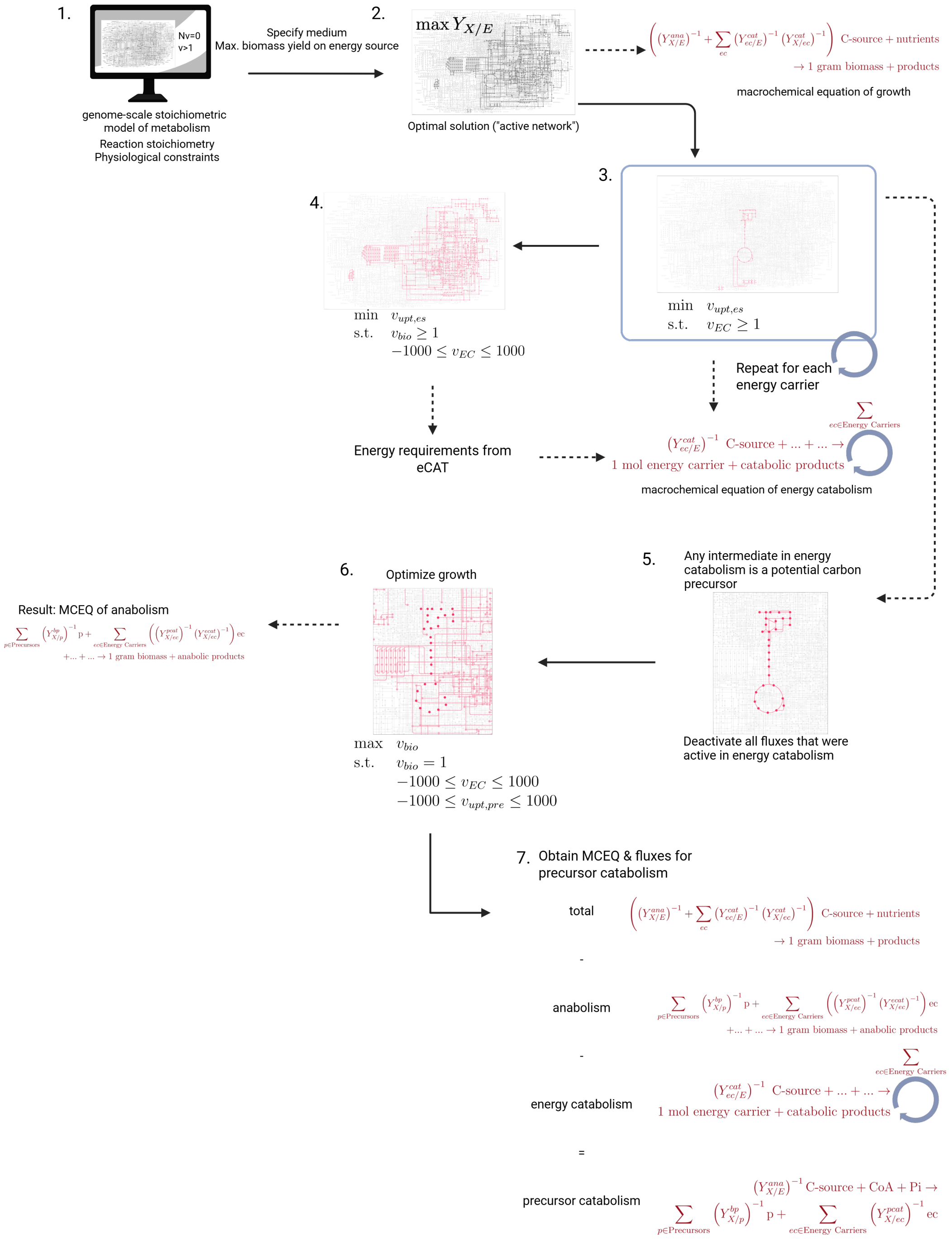

Figure S1: **Overview of methods to find the subnetworks of energy catabolism, precursor catabolism and anabolism from the active flux distribution after optimization.** The methods are described in Section A.

We have now determined the flux distributions to produce 1 (mmol) energy carrier, given the active metabolic network specification of the original problem. However, we do not yet know which energy carriers (ATP and NAPH or only ATP) are required by anabolism and how much we need of each them. This we determine from the anabolic flux distribution of $\boldsymbol{N}_A$. To find it, we add supply fluxes of each energy carrier present in the model (i.e. the reverse of e.g. Eqs. S5 and S6) to the active model $\boldsymbol{N}_A$. Just as some energy carriers cannot be produced independently of one another, they also cannot be consumed independently. As a result, there could be negative fluxes in anabolism, but only the reactions that were positive in catabolism (as the sum of both should equal the original biomass-optimal EFM, which by definition only has positive fluxes). In addition, we define that $CO_2$ cannot be shared between catabolism and anabolism. Thus, the lower bounds of all reactions that have a positive flux in catabolism, but do not have $CO_2$ as a by-product, are set to -1000. This is the reason that we first required a normalized flux distribution of catabolism, as before we could not define the bounds of anabolism. Then, the biomass objective function is set to 1 and the substrate uptake flux is minimized (Eq. S9, Figure S1, Step 4).

$$\begin{array}{lll} \text{minimize} & v_{upt,es} & \\ \text{subject to} & \boldsymbol{N}_A \cdot \boldsymbol{v} = \boldsymbol{0}, & \\ & -1000 \leq v_i \leq 1000 & \text{for } i > 0 \in v_{ec,norm} \text{ without } CO_2 \text{ as by-product,} \\ & 0 \leq v_i \leq 1000 & \text{for all other} i = 1, ..., n \\ & 0 \leq v_{ec,supply,j} \leq 1000 & \text{for all ec's j} \\ & v_{BOF} \geq 1 & \end{array} \tag{S9}$$

The optimal solution of this linear program is the anabolic flux vector of $\boldsymbol{N}_A$ we call $\boldsymbol{v}_{ANAec}^{opt}$.

We have now found an anabolic flux distribution that can contain any combination of the energy carriers present. There could be multiple combinations possible, i.e. energy can be exchanged between different energy carriers. In many organisms, for example, NADH can donate its electrons in oxidative phosphorylation, in which ATP is formed as a product. If ATP is defined as energy carrier in this case, potential required NADH can be produced in anabolism by running oxidative phosphorylation in reverse. Thus, any combination of NADH and ATP could be used as exchanged energy carrier. To determine which energy carriers are essential and which could replace each other, flux variability analysis (FVA) is performed within the optimal solution space of the anabolic model. If a supply flux of an energy carrier has a non-zero value without variability, the energy carrier is essential. If there are multiple energy carrier supply fluxes with variability, they might be interchangeable. Thus, the value of one energy carrier supply flux is set to its maximum, and another FVA is performed. If the values of the other energy carrier supply fluxes are zero without variability, they were interchangeable. Otherwise, the procedure is repeated until there is no variability in the energy carrier supply fluxes. The choice in order of energy carrier supply fluxes that is set to its maximal value partially determines the energy carriers that are exchanged. This procedure leads to the anabolic flux vector we continue with ($\boldsymbol{v}_{ANAec}$).

Then, FBA was performed in the anabolic model with only the required energy carriers. This way, the quantity of energy carriers is that is produced by catabolism is determined. Thus, the normalized energy carrier producing flux distributions that were determined before are multiplied with the value of their respective supply fluxes in anabolism (Eq. S10).

$$\boldsymbol{v_{ec}} = \boldsymbol{v_{ec,norm}} \cdot \frac{v_{ANAec}^{supply}}{v_{ec,norm}^{demand}} \tag{S10}$$

Then, all energy carrier producing flux distributions ($\boldsymbol{v_{ec,j}}$) are summed to form the catabolic flux distribution ($\boldsymbol{v_{eCAT}}$) (Eq. S11).

$$\boldsymbol{v_{eCAT}} = \sum_j \boldsymbol{v_{ec,j}} \tag{S11}$$

As verification, it was checked whether the catabolic and anabolic flux distribution summed to the original biomass-optimal flux distribution (Eq. S12).

$$v_A^{opt} = v_{ANAec} + v_{eCAT} \tag{S12}$$

#### A.3 Step 2: Dissection based on carbon precursors

Next, we aim to determine the flux distribution for anabolism from carbon precursors and energy carriers, instead of from substrate and energy carriers as defined before. For this, we start with the previously defined (energy-)catabolic and anabolic flux distributions. We can calculate the flux distribution for precursor catabolism, by subtracting the newly calculated anabolic flux distribution ($v_{ANA}$; Note that this is a different flux distribution than $v_{ANAec}$) and the previously calculated energy-catabolic flux distribution ($veCAT$) from the original total flux distribution ($v_A^{opt}$) (Eq. S13).

$$v_{pCAT} = v_A^{opt} - v_{eCAT} - v_{ANA} \tag{S13}$$

All metabolites in the catabolic flux distribution are potential carbon precursors, with the exception of compounds that do not contain carbon (such as $H_2O$ and $O_2$) and energy carriers. To find all energy carriers, the conserved moieties of the catabolic flux distribution were calculated by performing Gaussian elimination on its active stoichiometric matrix [2]. The conserved moieties related to the attachment of chemical groups, such as CoA, Pi and THF, were discarded and the rest were considered energy carriers.

Then, for each potential carbon precursors, exchange (e.g. Eq S14) and transfer (e.g. Eq S15) reactions were added to the model. The transfer reactions do not have any transport costs and do not include a periplasmic metabolite. If the carbon precursor has for example CoA or Pi attached, that group is released again in the exchange reaction (e.g. Eq S16), and a transfer reaction for the groups is additionally added. The exchange and transfer reactions are reversible to allow for anabolic byproducts. In addition, for each energy carrier that was previously determined, a chemically balanced supply reaction was added, like Eq. S5 and S6.

$$\text{pyruvate[ex]} \rightleftharpoons \tag{S14}$$

$$\text{pyruvate[ex]} \rightleftharpoons \text{pyruvate[c]} \tag{S15}$$

$$\text{acetyl}-\text{CoA[ex]} \rightleftharpoons \text{CoA[ex]} \tag{S16}$$

The resulting stoichiometric matrix is $\boldsymbol{N_{anaCP}}$.

The reactions in the catabolic flux distribution were deactivated by setting both the lower and upper bound to 0 (Figure S1, Step 5). This was done to prevent the consumption of the energetically most favorable precursor to form ATP. For example, the model could convert fructose-6-phosphate into 2 pyruvate and gain 3 ATP, instead of using the pyruvate from the exchange reaction.

To find a flux distribution for biomass production from carbon precursors, the BOF was set to 1 $h^{-1}$ and the BOF was optimized (Eq. S17, Figure S1, Step 6). This way, a flux distribution that allows growth using the precursors is found. There is no guarantee that this is the only flux distribution that is possible. However, as the current model stems from an EFM, with the only additions being the supplies of the carbon precursors and the energy carriers, the main variability is expected to be in the added reactions (See Section B.5).

$$\begin{array}{lll} \text{minimize} & v_{BOF} & \\ \text{subject to} & \boldsymbol{N_{anaCP}} \cdot \boldsymbol{v} = \boldsymbol{0}, & \\ & -1000 \leq v_i \leq 1000 & \text{for } i > 0 \in \text{exchange and transfer} \\ & 0 \leq v_i \leq 1000 & \text{for all other } i = 1, ..., n \\ & -1000 \leq v_{ec,supply,j} \leq 1000 & \text{for all ec's j} \\ & v_{BOF} = 1 & \end{array} \tag{S17}$$

The flux distribution for precursor catabolism is found by subtracting the anabolic and energy catabolic flux distributions from the total active flux distribution (Figure S1, Step 7).

#### A.4 Determining macrochemical equations

To obtain a clear overview of a process, the macrochemical equation (MCEQ) is calculated. From a genome scale model, this can be done by removing all exchange reactions from the (active) stoichiometric matrix (obtaining $N_{A,mceq}$) and the (active) flux vector (obtaining $v_{mceq}$). Then, the change in metabolites over time can be calculated using Eq. S18, in which $m$ is a vector of metabolite names.

$$\frac{d\boldsymbol{x_{ex}}}{dt} = \boldsymbol{m}^T \cdot \boldsymbol{N_{A,mceq}} \cdot \boldsymbol{v_{mceq}} \tag{S18}$$

In the case of an anabolic or catabolic flux distribution, the energy carrier supply and demand fluxes are removed from the stoichiometric matrix in addition to all the exchange fluxes. Then, the energy carriers that are consumed and produced appear in the MCEQ. The equations to calculate the MCEQ for the biomass-optimal flux distribution ($\frac{dx_{ex}^{opt}}{dt}$, Eq. S19) are given below.

$$\frac{d\boldsymbol{x_{ex}^{opt}}}{dt} = \boldsymbol{m}^T \cdot \boldsymbol{N_{A,mceq}} \cdot \boldsymbol{v_{mceq}^{opt}} \tag{S19}$$

with the (generalized) result, if the carbon source is also the energy source:

$$\begin{aligned} Y_{X/E}^{-1}\ \text{C-source} + n\ \text{N-source} + p\ \text{P-source} + s\ \text{S-source} + e\ \text{e-acceptor/donor} \\ \rightarrow 1\ \text{gram biomass} + b\ \text{byproducts} \end{aligned} \tag{S20}$$

This procedure can be repeated for all subnetworks to obtain their respective MCEQs. As an example, we give all MCEQs for *E. coli* growing aerobically on glucose in Section B.2.

#### A.5 Genome scale model adaptations

##### A.5.1 E. coli (iML1515)

The GAM and NGAM costs were adapted to be equal to those of the previous model iJO1366 [3]. The GAM was 53.95 $\text{mmol}_{ATP}$ $\text{gDW}^{-1}$ and the NGAM was 3.15 $\text{mmol}_{ATP}$ $\text{gDW}^{-1}$ $\text{h}^{-1}$.

The BOF of iML1515 [4] contained all biomass precursors, such as amino acids and (deoxy-)nucleotides separately. All ATP costs were lumped into one growth-associated maintenance (GAM) cost. To determine the energetic costs of each macromolecule, the BOF was rewritten to the following form (Eq. S21). The total GAM costs were distributed over each macromolecule, assuming the ATP costs as described in Fuchs (1998) [5] (Table S1). In addition, the tRNA loading of each molecule was modeled explicitly, similar to the genome scale model of *S. cerevisiae* [6].

$$\begin{aligned} &- 1.0\,\text{protein} - 1.0\,\text{lipid} - 1.0\,\text{lps} - 1.0\,\text{murein} - 1.0\,\text{DNA} - 1.0\,\text{RNA} \\ &- 1.0\,\text{cofactor} - 1.0\,\text{ion} - 16.12\,\text{ATP} - 16.12\,\text{H}_2\text{O} + 1.0\,\text{biomass} + 16.12\,\text{ADP} + 16.12\,\text{Pi} + 16.12\,\text{H}^+ \end{aligned} \tag{S21}$$

##### A.5.2 S. cerevisiae (Yeast9)

Yeast9 [6] is a consensus model for yeast. The reaction and metabolite ids were adapted to be human-readable to facilitate interpretation of results. For this, the dictionaries supplied on the Yeast-GEM GitHub were used. In addition, the stoichiometry of the mitochondrial ATP synthase was adapted to utilize four protons (Eq. S22) instead of three to reduce the P/O ratio and approach experimentally determined values [7].

$$-1.0\,\text{ADP} - 4.0\,\text{H}^+[\text{c}] - 1.0\,\text{Pi} + 1.0\,\text{ATP} + 3.0\,\text{H}^+[\text{m}] + 1.0\,\text{H}_2\text{O} \tag{S22}$$

The biomass objective function of Yeast9 was already formulated with macromolecules, but the GAM cost was lumped. Thus, the same costs were assumed for the biosynthesis of macromolecules from their building blocks as *E. coli*, but they were corrected for the macromolecular composition of *S. cerevisiae* [8, 9] (Table S1).

Table S1: Biomass composition of *E. coli* and *S. cerevisiae* from [8, 9] and ATP costs of polymerization of macromolecular building blocks from [5]. The protein costs were only the costs for initiation and elongation, as the tRNA loading is modelled explicitly. The polysaccharide elongation cost was set to zero as the polysaccharides are explicitly made in the yeast model.

| | *E. coli* biomass composition (g $gDW^{-1}$) | *E. coli* ATP cost ($mmol_{ATP}$ $gDW^{-1}$) | *S. cerevisiae* biomass composition (g $gDW^{-1}$) | *S. cerevisiae* ATP cost ($mmol_{ATP}$ $gDW^{-1}$) |
|---|---|---|---|---|
| Protein | 0.68 | 21.4 | 0.51 | 16.9 |
| Lipid | 0.15 | 0.3 | 0.07 | 0.144 |
| Lipopolysaccharide | 0.01 | 0 | 0 | - |
| DNA | 0.011 | 0.3 | 0.01 | 0.27 |
| RNA | 0.07 | 1.5 | 0.11 | 2.7 |
| Polysaccharides | 0.002 | 0.2 | 0.27 | 0 |
| Cofactor dilution | 0 | 0 | 0 | 0 |
| Ion dilution | 0 | 0 | 0 | 0 |
| GAM | | 16.12 | | 35.7 |

##### A.5.3 Synechocystis (iSynCJ816)

The *Synechocystis* GSMM requires some adaptations for autotrophic growth, as described in Joshi et al. (2017). Additionally, the reaction 'R_HCO3E1' produced internal $CO_2$ from nothing instead of being equilibrium between $CO_2$ and $HCO_3^-$. Thus, $HCO_3^-$ was added to the reaction as a substrate. Lastly, the reaction 'R_CAT' enabled futile oxygen consumption and was thus shut down (i.e. the lower and upper bound were set to 0).

### B Supplementary results

#### B.1 List of simulations

Table S2: Overview of performed simulations and the exchanged energy carriers.

| Organism | Energy source | Additional substrates | Genome scale model | Exchanged energy carriers |
|---|---|---|---|---|
| *C. ljungdahlii* | $H_2$ | $CO_2$, $NH_4^+$, $SO_4^{2-}$, $PO_4^{3-}$ | iHN637 [11] | Fd |
| *C. ljungdahlii* | CO | $NH_4^+$, $SO_4^{2-}$, $PO_4^{3-}$ | iHN637 [11] | ATP, Fd |
| *C. ljungdahlii* | CO | $NH_4^+$, $SO_4^{2-}$, $PO_4^{3-}$ | iHN637 [11] | ATP |
| *G. metallireducens* | acetate | $Fe_3^+$, $NH_4^+$, $SO_3^{2-}$, $SO_4^{2-}$, $PO_4^{3-}$ | iAF987 [12] | ATP, NADPH |
| *G. metallireducens* | acetate | $NO_3^-$, $SO_3^{2-}$, $SO_4^{2-}$, $PO_4^{3-}$ | iAF987 [12] | ATP, NADPH |
| *G. metallireducens* | butyrate | $Fe_3^+$, $NH_4^+$, $SO_3^{2-}$, $SO_4^{2-}$, $PO_4^{3-}$ | iAF987 [12] | ATP, NADPH |
| *G. metallireducens* | ethanol | $Fe_3^+$, $NH_4^+$, $SO_3^{2-}$, $SO_4^{2-}$, $PO_4^{3-}$ | iAF987 [12] | ATP, NADPH |
| *G. metallireducens* | ethanol | $NO_2^-$, $SO_3^{2-}$, $SO_4^{2-}$, $PO_4^{3-}$ | iAF987 [12] | ATP |
| *G. metallireducens* | formate | $Fe_3^+$, $NH_4^+$, $SO_3^{2-}$, $SO_4^{2-}$, $PO_4^{3-}$ | iAF987 [12] | ATP, NADH |
| *M. barkeri* | acetate | $NH_4^+$, cysteine, $PO_4^{3-}$ | iMG746 [13] | ATP |
| *M. barkeri* | $H_2$ | $HCO_3^-$, $NH_4^+$, cysteine, $PO_4^{3-}$ | iMG746 [13] | ATP, Fd |
| *S. cerevisiae* | glucose | $O_2$, $NH_4^+$, $SO_4^{2-}$, $PO_4^{3-}$ | Yeast9 [6] | ATP, NADPH |
| *S. cerevisiae* | galactose | $O_2$, $NH_4^+$, $SO_4^{2-}$, $PO_4^{3-}$ | Yeast9 [6] | ATP, NADPH |
| *S. cerevisiae* | lactate | $O_2$, $NH_4^+$, $SO_4^{2-}$, $PO_4^{3-}$ | Yeast9 [6] | ATP, NADPH |
| *S. cerevisiae* | maltotriose | $O_2$, $NH_4^+$, $SO_4^{2-}$, $PO_4^{3-}$ | Yeast9 [6] | ATP, NADPH |
| *S. cerevisiae* | pyruvate | $O_2$, $NH_4^+$, $SO_4^{2-}$, $PO_4^{3-}$ | Yeast9 [6] | ATP, NADPH |
| *S. cerevisiae* | ribose | $O_2$, $NH_4^+$, $SO_4^{2-}$, $PO_4^{3-}$ | Yeast9 [6] | ATP, NADPH |
| *P. putida* | acetate | $O_2$, $NH_4^+$, $SO_4^{2-}$, $PO_4^{3-}$ | iJN1462 [14] | ATP |
| *P. putida* | ethanol | $O_2$, $NH_4^+$, $SO_4^{2-}$, $PO_4^{3-}$ | iJN1462 [14] | ATP |
| *P. putida* | glucose | $O_2$, $NH_4^+$, $SO_4^{2-}$, $PO_4^{3-}$ | iJN1462 [14] | ATP |
| *P. putida* | glycerol | $O_2$, $NH_4^+$, $SO_4^{2-}$, $PO_4^{3-}$ | iJN1462 [14] | ATP |
| *P. putida* | hexadecanoate 16:1 | $O_2$, $NH_4^+$, $SO_4^{2-}$, $PO_4^{3-}$ | iJN1462 [14] | ATP |
| *P. putida* | lactate | $O_2$, $NH_4^+$, $SO_4^{2-}$, $PO_4^{3-}$ | iJN1462 [14] | NADH |

| | | | | |
|---|---|---|---|---|
| *P. putida* | methanol | $O_2$, $NH_4^+$, $SO_4^{2-}$, $PO_4^{3-}$ | iJN1462 [14] | ATP |
| *P. putida* | octadecanoate | $O_2$, $NH_4^+$, $SO_4^{2-}$, $PO_4^{3-}$ | iJN1462 [14] | ATP |
| *P. putida* | putrescine | $O_2$, $NH_4^+$, $SO_4^{2-}$, $PO_4^{3-}$ | iJN1462 [14] | ATP |
| *E. coli* | acetate | $O_2$, $NH_4^+$, $SO_4^{2-}$, $PO_4^{3-}$ | iML1515 [4] | ATP |
| *E. coli* | $\alpha$-ketoglutarate | $O_2$, $NH_4^+$, $SO_4^{2-}$, $PO_4^{3-}$ | iML1515 [4] | ATP |
| *E. coli* | citrate | $O_2$, $NH_4^+$, $SO_4^{2-}$, $PO_4^{3-}$ | iML1515 [4] | ATP, NADPH |
| *E. coli* | fructose | $O_2$, $NH_4^+$, $SO_4^{2-}$, $PO_4^{3-}$ | iML1515 [4] | ATP, NADPH |
| *E. coli* | fumarate | $O_2$, $NH_4^+$, $SO_4^{2-}$, $PO_4^{3-}$ | iML1515 [4] | ATP, NADPH |
| *E. coli* | galactose | $O_2$, $NH_4^+$, $SO_4^{2-}$, $PO_4^{3-}$ | iML1515 [4] | ATP, NADPH |
| *E. coli* | gluconate | $O_2$, $NH_4^+$, $SO_4^{2-}$, $PO_4^{3-}$ | iML1515 [4] | ATP, NADPH |
| *E. coli* | glucose | $O_2$, $NH_4^+$, $SO_4^{2-}$, $PO_4^{3-}$ | iML1515 [4] | ATP, NADPH |
| *E. coli* | glucose | $NH_4^+$, $SO_4^{2-}$, $PO_4^{3-}$ | iML1515 [4] | ATP, NADH |
| *E. coli* | glycerol | $O_2$, $NH_4^+$, $SO_4^{2-}$, $PO_4^{3-}$ | iML1515 [4] | ATP |
| *E. coli* | lactate | $O_2$, $NH_4^+$, $SO_4^{2-}$, $PO_4^{3-}$ | iML1515 [4] | ATP |
| *E. coli* | malate | $O_2$, $NH_4^+$, $SO_4^{2-}$, $PO_4^{3-}$ | iML1515 [4] | ATP, NADPH |
| *E. coli* | pyruvate | $O_2$, $NH_4^+$, $SO_4^{2-}$, $PO_4^{3-}$ | iML1515 [4] | ATP |
| *E. coli* | ribose | $O_2$, $NH_4^+$, $SO_4^{2-}$, $PO_4^{3-}$ | iML1515 [4] | ATP, NADPH |
| *E. coli* | sorbitol | $O_2$, $NH_4^+$, $SO_4^{2-}$, $PO_4^{3-}$ | iML1515 [4] | ATP, NADPH |
| *E. coli* | succinate | $O_2$, $NH_4^+$, $SO_4^{2-}$, $PO_4^{3-}$ | iML1515 [4] | ATP, NADPH |
| *E. coli* | xylose | $O_2$, $NH_4^+$, $SO_4^{2-}$, $PO_4^{3-}$ | iML1515 [4] | ATP, NADPH |
| *E. coli* | alanine | $O_2$, $NH_4^+$, $SO_4^{2-}$, $PO_4^{3-}$ | iML1515 [4] | ATP |
| *E. coli* | arginine | $O_2$, $NH_4^+$, $SO_4^{2-}$, $PO_4^{3-}$ | iML1515 [4] | ATP |
| *E. coli* | aspartate | $O_2$, $NH_4^+$, $SO_4^{2-}$, $PO_4^{3-}$ | iML1515 [4] | ATP |
| *E. coli* | glutamate | $O_2$, $NH_4^+$, $SO_4^{2-}$, $PO_4^{3-}$ | iML1515 [4] | ATP |
| *E. coli* | glutamine | $O_2$, $NH_4^+$, $SO_4^{2-}$, $PO_4^{3-}$ | iML1515 [4] | ATP |
| *E. coli* | tryptophan | $O_2$, $NH_4^+$, $SO_4^{2-}$, $PO_4^{3-}$ | iML1515 [4] | ATP |

| | | | | |
|---|---|---|---|---|
| *Synechocystis* | photons | $HCO_3^-$, $NO_3^-$, $SO_4^{2-}$, $PO_4^{3-}$ | iSynCJ816 [10] | NADH |

#### B.2 Example simulation with associated macrochemical equations: E. coli growing on aerobically on glucose

As an example, we present all macrochemical equations for the dissection of the metabolism of *E. coli* growing aerobically on glucose into energy catabolism, precursor catabolism and anabolism. After that, we show the energy and substrate requirements for the macromolecules.

We start with the overall MCEQ, that follows from minimization of glucose uptake, while the BOF is fixed. This is the macrochemical equation with the optimal biomass yield on substrate. The units of the metabolites in the equations are in mmol, except for biomass, which is in gram. For the macromolecules, the unit is the number of grams of biomass which can be generated using this macromolecule, e.g. for a value of 1, 1 gram biomass can be made per this amount of protein.

**Growth**

$$\begin{aligned}-2\cdot10^{-06}\ \text{biotin}-0.0052\ \text{Ca}^{2+}-0.0052\ \text{Cl}^{-}-2.5\cdot10^{-05}\ \text{Co}^{2+}-0.00071\ \text{Cu}^{2+}\\ -0.016\ \text{Fe}^{2+}-10.00\ \text{glucose}-0.2\ \text{K}^{+}-0.0087\ \text{Mg}^{2+}-0.00069\ \text{Mn}^{2+}\\ -7\cdot10^{-06}\ \text{MoO}_4^{2-}-10.80\ \text{NH}_4^{+}-0.00032\ \text{Ni}^{2+}-16.82\ \text{O}_2-0.96\ \text{PO}_4^{3-}\\ -0.25\ \text{SO}_4^{2-}-0.00034\ \text{Zn}^{2+}+1.00\ \text{gram biomass}+18.96\ \text{CO}_2+45.36\ \text{H}_2\text{O}+9.19\ \text{H}^{+}\end{aligned} \tag{S23}$$

After obtaining the flux distribution with the optimal $Y_{X/E}$, all inactive reactions are deleted to obtain the active model. To obtain the energy conserving flux distribution, we add an ATP hydrolysis reaction, set the lower and upper bound to 1. In addition, the constraint on biomass production is removed. Then, the glucose uptake flux is minimized. As the resulting MCEQ contained biomass, the next step was to correct for the biomass. This was all explained in the Appendix section A.4 above. This leads to the flux distribution and corresponding MCEQ for the production of 1 ATP. In this flux distribution, negative fluxes are present in the oxidative pentose phosphate pathway which consume the NADPH that is generated in the TCA cycle.

**ATP production (not corrected for biomass)**

$$\begin{aligned}-1.00\ \text{ADP}-9.9\cdot10^{-05}\ \text{Ca}^{2+}-9.9\cdot10^{-05}\ \text{Cl}^{-}-4.7\cdot10^{-07}\ \text{Co}^{2+}\\ -1.3\cdot10^{-05}\ \text{Cu}^{2+}-0.0003\ \text{Fe}^{2+}-0.23\ \text{glucose}-0.0037\ \text{K}^{+}-0.00016\ \text{Mg}^{2+}\\ -1.3\cdot10^{-05}\ \text{Mn}^{2+}-1.3\cdot10^{-07}\ \text{MoO}_4^{2-}-0.21\ \text{NH}_4^{+}-6.1\cdot10^{-06}\ \text{Ni}^{2+}-0.57\ \text{O}_2\\ -0.018\ \text{PO}_4^{3-}-0.0048\ \text{SO}_4^{2-}-6.5\cdot10^{-06}\ \text{Zn}^{2+}+1.00\ \text{ATP}+0.019\ \text{gram biomass}\\ +0.61\ \text{CO}_2+1.11\ \text{H}_2\text{O}+0.17\ \text{H}^{+}\end{aligned} \tag{S24}$$

**ATP production (corrected for biomass)**

$$\begin{aligned}-1\ \text{ADP}-0.042\ \text{glucose}-0.25\ \text{O}_2+1\ \text{ATP}+0.25\ \text{CO}_2\\ +0.25\ \text{H}_2\text{O}\end{aligned} \tag{S25}$$

The same process is repeated for each energy carrier present in the genome scale model. In this case, it later turns out that only NADPH is required as input for anabolism. In the flux distribution for the production of NADPH, glucose-6-phosphate isomerase (PGI) has a negative flux, to be able to fully convert glucose into $CO_2$ via the oxPPP. The required amounts of energy carriers are determined when calculating the anabolic flux distribution.

**NADPH production (not corrected for biomass)**

$$\begin{aligned}-1.7\cdot10^{-07}\ \text{biotin}-0.00045\ \text{Ca}^{2+}-0.00045\ \text{Cl}^{-}-2.1\cdot10^{-06}\ \text{Co}^{2+}\\ -6.1\cdot10^{-05}\ \text{Cu}^{2+}-0.0014\ \text{Fe}^{2+}-0.96\ \text{glucose}-0.017\ \text{K}^{+}-0.00074\ \text{Mg}^{2+}\\ -5.9\cdot10^{-05}\ \text{Mn}^{2+}-6\cdot10^{-07}\ \text{MoO}_4^{2-}-1.00\ \text{NADP}-0.93\ \text{NH}_4^{+}\\ -2.8\cdot10^{-05}\ \text{Ni}^{2+}-1.57\ \text{O}_2-0.083\ \text{PO}_4^{3-}-0.022\ \text{SO}_4^{2-}-2.9\cdot10^{-05}\ \text{Zn}^{2+}\\ +0.086\ \text{gram biomass}+2.25\ \text{CO}_2+3.51\ \text{H}_2\text{O}+2.79\ \text{H}^{+}+1.00\ \text{NADPH}\end{aligned} \tag{S26}$$

**NADPH production (corrected for biomass)**

$$-0.1 \text{ glucose} - 0.37 \text{ H}_2\text{O} - 1.00 \text{ NADP} - 0.13 \text{ O}_2 + 0.63 \text{ CO}_2 + 2.00 \text{ H}^+ + 1.00 \text{ NADPH} \quad (S27)$$

To find the anabolic flux distribution, the linear program in Equation S9 is solved. Then, only the supplies of the required energy carriers carry flux.

**Anabolism (energy carriers)**

$$\begin{aligned} -28.33 \text{ ATP} - 2 \cdot 10^{-06} \text{ biotin} - 0.0052 \text{ Ca}^{2+} - 0.0052 \text{ Cl}^{-} - 2.5 \cdot 10^{-05} \text{ Co}^{2+} \\ -0.00071 \text{ Cu}^{2+} - 0.016 \text{ Fe}^{2+} - 7.55 \text{ glucose} - 15.17 \text{ H}^{+} - 0.2 \text{ K}^{+} - 0.0087 \text{ Mg}^{2+} \\ -0.00069 \text{ Mn}^{2+} - 7 \cdot 10^{-06} \text{ MoO}_4^{2-} - 12.18 \text{ NADPH} - 10.80 \text{ NH}_4^{+} - 0.00032 \text{ Ni}^{2+} \\ -8.22 \text{ O}_2 - 0.96 \text{ PO}_4^{3-} - 0.25 \text{ SO}_4^{2-} - 0.00034 \text{ Zn}^{2+} + 28.33 \text{ ADP} + 1 \text{ gram biomass} \\ +4.26 \text{ CO}_2 + 42.85 \text{ H}_2\text{O} + 12.18 \text{ NADP} \end{aligned} \quad (S28)$$

The energy carrier production flux distributions are multiplied with the required amount from the anabolic flux distribution. Then, they are summed to find the energy conserving flux distribution, which concludes the dissection of growth into energy catabolism and anabolism.

**ATP production (scaled to biomass)**

$$\begin{aligned} -28.33 \text{ ADP} - 1.18 \text{ glucose} - 7.08 \text{ O}_2 + 28.33 \text{ ATP} + 1.1 \cdot 10^{-06} \text{ biotin} + 7.08 \text{ CO}_2 \\ +7.08 \text{ H}_2\text{O} \end{aligned} \quad (S29)$$

**NADPH production (scaled to biomass)**

$$-1.27 \text{ glucose} - 4.57 \text{ H}_2\text{O} - 12.18 \text{ NADP} - 1.52 \text{ O}_2 + 7.61 \text{ CO}_2 + 24.36 \text{ H}^+ + 12.18 \text{ NADPH} \quad (S30)$$

**Energy catabolism**

$$\begin{aligned} -28.33 \text{ ADP} - 2.45 \text{ glucose} - 12.18 \text{ NADP} - 8.61 \text{ O}_2 + 28.33 \text{ ATP} \\ +14.69 \text{ CO}_2 + 2.52 \text{ H}_2\text{O} + 24.36 \text{ H}^+ + 12.18 \text{ NADPH} \end{aligned} \quad (S31)$$

Then, all reactions in the subnetwork of energy catabolism are deactivated, and all intermediate metabolites are supplied. To simulate the production of 1 gram of biomass, the lower and upper bound of the BOF are set to 1.0 and BOF is optimized. This way, a (non-unique) solution is obtained for biomass production.

**Anabolism (precursors):**

$$\begin{aligned} -1.72 \text{ 3P-glycerate} - 3.17 \text{ acetyl-CoA} - 1.08\ \alpha\text{-ketoglutarate} - 65.17 \text{ ATP} \\ -2 \cdot 10^{-06} \text{ biotin} - 0.0052 \text{ Ca}^{2+} - 0.0052 \text{ Cl}^{-} - 1.02 \text{ CO}_2 - 2.5 \cdot 10^{-05} \text{ Co}^{2+} \\ -0.00071 \text{ Cu}^{2+} - 0.14 \text{ dihydroxyacetone-P} - 0.38 \text{ erythrose-4P} - 0.094 \text{ fructose-6P} - 0.016 \text{ Fe}^{2+} \\ -29.04 \text{ H}^{+} - 0.2 \text{ K}^{+} - 0.0087 \text{ Mg}^{2+} - 0.00069 \text{ Mn}^{2+} - 7 \cdot 10^{-06} \text{ MoO}_4^{2-} - 1.89 \text{ NADH} \\ -13.25 \text{ NADPH} - 10.80 \text{ NH}_4^{+} - 0.00032 \text{ Ni}^{2+} - 0.17 \text{ O}_2 - 3.75 \text{ phosphoenol-P} - 2.95 \text{ pyruvate} \\ -0.93 \text{ ribose-5P} - 0.04 \text{ ribulose-5P} - 0.25 \text{ SO}_4^{2-} - 0.52 \text{ succinyl-CoA} - 0.00034 \text{ Zn}^{2+} \\ +65.17 \text{ ADP} + 1.00 \text{ gram biomass} + 3.69 \text{ CoA} + 1.04 \text{ fumarate} + 0.054 \text{ glyceraldehyde-3P} \\ +23.04 \text{ H}_2\text{O} + 1.89 \text{ NAD}^{+} + 13.25 \text{ NADP} + 6.04 \text{ PO}_4^{3-} + 0.53 \text{ succinate} \end{aligned} \quad (S32)$$

Next, we obtained the MCEQ for precursor catabolism, by subtracting energy catabolism and biomass production from the total MCEQ.

$$\text{precursor production} = \text{total} - \text{biomass production} - \text{energy catabolism} \quad (S33)$$

**Precursor catabolism**

$$-36.84\ \text{ADP} - 3.69\ \text{CoA} - 1.04\ \text{fumarate} - 0.054\ \text{glyceraldehyde-3P} - 7.55\ \text{glucose} - 1.89\ \text{NAD}^+$$
$$-1.07\ \text{NADP} - 0.0047\ \text{NH}_4^+ - 8.05\ \text{O}_2 - 7.01\ \text{PO}_4^{3-} - 0.00067\ \text{SO}_4^{2-} - 0.53\ \text{succinate}$$
$$+1.72\ \text{3P-glycerate} + 3.17\ \text{acetyl-CoA} + 1.08\ \alpha\text{-ketoglutarate} + 36.84\ \text{ATP} + 5.28\ \text{CO}_2$$
$$+0.14\ \text{dihydroxyacetone-P} + 0.38\ \text{erythrose-4P} + 0.094\ \text{fructose-6P} + 19.81\ \text{H}_2\text{O} + 13.87\ \text{H}^+ + 1.89\ \text{NADH}$$
$$+1.07\ \text{NADPH} + 3.75\ \text{phosphoenol-P} + 2.95\ \text{pyruvate} + 0.93\ \text{ribose-5P} + 0.04\ \text{ribulose-5P}$$
$$+0.52\ \text{succinyl-CoA} \tag{S34}$$

Alternatively, we could have optimized the production of each precursor separately. This gives equivalent results, as we have seen from the dissection into macromolecular subnetworks, but is more laborious.

##### B.2.1 Macromolecule production in the active network

Next, the metabolic network can be dissected into the production of different macromolecules if the BOF is formulated in terms of macromolecules. The networks of macromolecule production are in turn dissected into energy catabolism and anabolic networks. We do not show the precursor requirements for each macromolecule, and the anabolism in this case is thus the anabolism from precursors together with precursor catabolism.

We start with protein. The production of protein was optimized. Similar to the MCEQ of ATP production, this MCEQ also produces biomass. This is due to the fact that the production networks of different macromolecules share resources, such as energy carriers.

**Protein (uncorrected for biomass)**

$$-6.7\cdot10^{-06}\ \text{biotin} - 0.017\ \text{Ca}^{2+} - 0.017\ \text{Cl}^- - 8.4\cdot10^{-05}\ \text{Co}^{2+} - 0.0024\ \text{Cu}^{2+}$$
$$-0.054\ \text{Fe}^{2+} - 39.72\ \text{glucose} - 0.66\ \text{K}^+ - 0.029\ \text{Mg}^{2+} - 0.0023\ \text{Mn}^{2+}$$
$$-2.4\cdot10^{-05}\ \text{MoO}_4^{2-} - 43.61\ \text{NH}_4^+ - 0.0011\ \text{Ni}^{2+} - 66.24\ \text{O}_2 - 3.24\ \text{PO}_4^{3-}$$
$$-1.09\ \text{SO}_4^{2-} - 0.0011\ \text{Zn}^{2+} + 1.00\ \text{protein} + 3.36\ \text{gram biomass} + 74.80\ \text{CO}_2 + 179.88\ \text{H}_2\text{O}$$
$$+37.57\ \text{H}^+ \tag{S35}$$

The MCEQ for protein is corrected for biomass production. Now we can observe that the production of protein consumes about 61% of the total glucose required for growth.

**Protein**

$$-6.12\ \text{glucose} - 7.33\ \text{NH}_4^+ - 9.72\ \text{O}_2 - 0.25\ \text{SO}_4^{2-} + 1.00\ \text{protein} + 11.11\ \text{CO}_2 + 27.48\ \text{H}_2\text{O} + 6.70\ \text{H}^+ \tag{S36}$$

To dissect the production of protein into an energy catabolic and anabolic network, the same steps are repeated as with dissection of growth: an energy carrier flux is optimized and the flux distribution is corrected for biomass production if necessary. In addition to biomass production, there might be protein production in the flux distribution for an energy carrier, thus the flux distribution is corrected if required. As expected, the flux distributions for the production of energy carriers while producing a macromolecule and biomass were the same. The only thing that changed was the relative contributions of the ATP and NADPH producing flux distributions to energy catabolism.

**Protein energy catabolism**

$$-10.16\ \text{ADP} - 1.55\ \text{glucose} - 1.51\ \text{H}_2\text{O} - 10.79\ \text{NADP} - 3.89\ \text{O}_2 + 10.16\ \text{ATP} + 9.28\ \text{CO}_2$$
$$+21.58\ \text{H}^+ + 10.79\ \text{NADPH} \tag{S37}$$

In the anabolic macrochemical equation for protein production, biomass appeared again. In this case, the correction for biomass production is performed by subtracting the anabolic network for growth (with the appropriate factor) from the obtained flux distribution. This is required because the biomass-producing part of the obtained flux distribution consumed ATP and NADPH, i.e. the anabolic network was active, not the catabolic network. The resulting anabolic MCEQ shows that a substantial fraction of NADPH (88.5%) that is required for growth, is required for the production of protein. In addition, while 61.2% of the glucose is used for production of protein, only 35.2% of ATP is required for protein production. This indicates that the anabolic ATP fraction of protein production is higher than average.

**Protein anabolism (not corrected for biomass)**

$$\begin{aligned} &-105.34\ \text{ATP} - 6.7\cdot 10^{-06}\ \text{biotin} - 0.017\ \text{Ca}^{2+} - 0.017\ \text{Cl}^{-} - 8.4\cdot 10^{-05}\ \text{Co}^{2+} \\ &-0.0024\ \text{Cu}^{2+} - 0.054\ \text{Fe}^{2+} - 29.95\ \text{glucose} - 65.84\ \text{H}^{+} - 0.66\ \text{K}^{+} - 0.029\ \text{Mg}^{2+} \\ &-0.0023\ \text{Mn}^{2+} - 2.4\cdot 10^{-05}\ \text{MoO}_4^{2-} - 51.71\ \text{NADPH} - 43.61\ \text{NH}_4^{+} - 0.0011\ \text{Ni}^{2+} \\ &-33.44\ \text{O}_2 - 3.24\ \text{PO}_4^{3-} - 1.09\ \text{SO}_4^{2-} - 0.0011\ \text{Zn}^{2+} + 1.00\ \text{protein} + 105.34\ \text{ADP} \\ &+3.36\ \text{gram biomass} + 16.15\ \text{CO}_2 + 172.94\ \text{H}_2\text{O} + 51.71\ \text{NADP} \end{aligned} \tag{S38}$$

**Protein anabolism**

$$\begin{aligned} &-10.16\ \text{ATP} - 4.58\ \text{glucose} - 14.88\ \text{H}^{+} - 10.79\ \text{NADPH} - 7.33\ \text{NH}_4^{+} - 5.84\ \text{O}_2 \\ &-0.25\ \text{SO}_4^{2-} + 1.00\ \text{protein} + 10.16\ \text{ADP} + 1.83\ \text{CO}_2 + 28.99\ \text{H}_2\text{O} + 10.79\ \text{NADP} \end{aligned} \tag{S39}$$

Using the same steps, the production networks of all macromolecules can be dissected. To avoid repetitiveness, only the total, energy-catabolic and anabolic MCEQs of the remaining macromolecules are shown. For the production of lipids, 11.7% of the glucose is required. Surprisingly, ATP and NADPH are byproducts of the anabolism of lipids. As a consequence, ATP and NADPH are substrates in the MCEQ for energy catabolism for lipids. Because the network must be balanced, glucose is now a product of this energy catabolism. The negative fluxes that the energy catabolism network have, are used to compensate ATP and NADPH consumption in other macromolecular subnetworks.

**Lipid**

$$\begin{aligned} &-1.17\ \text{glucose} - 0.14\ \text{H}^{+} - 0.14\ \text{NH}_4^{+} - 0.14\ \text{PO}_4^{3-} + 1.00\ \text{lipid} + 1.87\ \text{CO}_2 \\ &+2.44\ \text{H}_2\text{O} + 0.14\ \text{O}_2 \end{aligned} \tag{S40}$$

**Lipid energy catabolism**

$$\begin{aligned} &-2.09\ \text{ATP} - 1.08\ \text{CO}_2 - 0.19\ \text{H}_2\text{O} - 1.79\ \text{H}^{+} - 0.9\ \text{NADPH} + 2.09\ \text{ADP} + 0.18\ \text{glucose} \\ &+0.9\ \text{NADP} + 0.63\ \text{O}_2 \end{aligned} \tag{S41}$$

**Lipid anabolism**

$$\begin{aligned} &-2.09\ \text{ADP} - 1.35\ \text{glucose} - 0.9\ \text{NADP} - 0.14\ \text{NH}_4^{+} - 0.49\ \text{O}_2 - 0.14\ \text{PO}_4^{3-} \\ &+1.00\ \text{lipid} + 2.09\ \text{ATP} + 2.95\ \text{CO}_2 + 2.62\ \text{H}_2\text{O} + 1.66\ \text{H}^{+} + 0.9\ \text{NADPH} \end{aligned} \tag{S42}$$

Lipopolysaccharide synthesis consumes 3.5% of the glucose and has ATP and ATP as byproducts of its anabolism, similar to lipid synthesis.

**Lipopolysaccharide**

$$\begin{aligned} &-0.35\ \text{glucose} - 0.039\ \text{NH}_4^{+} - 0.039\ \text{O}_2 - 0.039\ \text{PO}_4^{3-} + 1.00\ \text{lipopolysaccharide} + 0.45\ \text{CO}_2 \\ &+0.7\ \text{H}_2\text{O} + 0.078\ \text{H}^{+} \end{aligned} \tag{S43}$$

**Lipopolysaccharide energy catabolism**

$$\begin{aligned} &-0.45\ \text{ATP} - 0.23\ \text{CO}_2 - 0.04\ \text{H}_2\text{O} - 0.39\ \text{H}^{+} - 0.19\ \text{NADPH} + 0.45\ \text{ADP} + 0.039\ \text{glucose} \\ &+0.19\ \text{NADP} + 0.14\ \text{O}_2 \end{aligned} \tag{S44}$$

**Lipopolysaccharide anabolism**

$$\begin{aligned}&-0.45\ \text{ADP} - 0.39\ \text{glucose} - 0.19\ \text{NADP} - 0.039\ \text{NH}_4^+ - 0.18\ \text{O}_2 - 0.039\ \text{PO}_4^{3-} \\ &+1.00\ \text{lipopolysaccharide} + 0.45\ \text{ATP} + 0.68\ \text{CO}_2 + 0.74\ \text{H}_2\text{O} + 0.46\ \text{H}^+ + 0.19\ \text{NADPH}\end{aligned} \tag{S45}$$

The production of RNA consumes 12.4% of the glucose. RNA is the only macromolecule that does not require NADPH from the energy conserving part for its growth. This can be explained from the precursor metabolites that RNA is made from: they are intermediates of the pentose phosphate pathway, which generates NADPH. So, the NADPH that is required in the formation of RNA is made in the pathway from glucose to RNA, and is thus not required from energy catabolism.

**RNA**

$$\begin{aligned}&-1.24\ \text{glucose} - 2.64\ \text{NH}_4^+ - 2.57\ \text{O}_2 - 0.67\ \text{PO}_4^{3-} + 1.00\ \text{RNA} + 1.07\ \text{CO}_2 + 8.49\ \text{H}_2\text{O} \\ &+1.97\ \text{H}^+\end{aligned} \tag{S46}$$

**RNA energy catabolism**

$$-4.27\ \text{ADP} - 0.18\ \text{glucose} - 1.07\ \text{O}_2 + 4.27\ \text{ATP} + 1.07\ \text{CO}_2 + 1.07\ \text{H}_2\text{O} \tag{S47}$$

**RNA anabolism**

$$\begin{aligned}&-4.27\ \text{ATP} - 1.07\ \text{glucose} - 2.64\ \text{NH}_4^+ - 1.50\ \text{O}_2 - 0.67\ \text{PO}_4^{3-} + 4.27\ \text{ADP} + 7.43\ \text{H}_2\text{O} \\ &+1.97\ \text{H}^+\end{aligned} \tag{S48}$$

2.2% of glucose is used for DNA synthesis. DNA synthesis requires only a small amount of ATP and NADPH, respectively 0.11% and 0.36% of the total from energy catabolism is used for DNA synthesis. This seems to be very little for such a vital process in the cell.

**DNA**

$$-0.22\ \text{glucose} - 0.4\ \text{NH}_4^+ - 0.42\ \text{O}_2 - 0.11\ \text{PO}_4^{3-} + 1.00\ \text{DNA} + 0.26\ \text{CO}_2 + 1.40\ \text{H}_2\text{O} + 0.29\ \text{H}^+ \tag{S49}$$

**DNA energy catabolism**

$$\begin{aligned}&-0.033\ \text{ATP} - 0.0032\ \text{glucose} - 0.025\ \text{H}_2\text{O} - 0.044\ \text{NADP} + 0.033\ \text{ADP} + 0.019\ \text{CO}_2 \\ &+0.088\ \text{H}^+ + 0.044\ \text{NADPH} + 0.0028\ \text{O}_2\end{aligned} \tag{S50}$$

**DNA anabolism**

$$\begin{aligned}&-0.033\ \text{ADP} - 0.21\ \text{glucose} - 0.044\ \text{NADPH} - 0.4\ \text{NH}_4^+ - 0.42\ \text{O}_2 - 0.11\ \text{PO}_4^{3-} \\ &+1.00\ \text{DNA} + 0.033\ \text{ATP} + 0.24\ \text{CO}_2 + 1.43\ \text{H}_2\text{O} + 0.21\ \text{H}^+ + 0.044\ \text{NADP}\end{aligned} \tag{S51}$$

The energy catabolism for murein synthesis has ATP and NADPH as a byproduct, similar to lipid an lipopolysaccharides.

**Murein**

$$-0.2\ \text{glucose} - 0.21\ \text{NH}_4^+ - 0.14\ \text{O}_2 + 1.00\ \text{murein} + 0.13\ \text{CO}_2 + 0.67\ \text{H}_2\text{O} + 0.26\ \text{H}^+ \tag{S52}$$

**Murein energy catabolism**

$$\begin{aligned}&-0.11\ \text{ATP} - 0.055\ \text{CO}_2 - 0.0095\ \text{H}_2\text{O} - 0.092\ \text{H}^+ - 0.046\ \text{NADPH} \\ &+0.11\ \text{ADP} + 0.0092\ \text{glucose} + 0.046\ \text{NADP} + 0.032\ \text{O}_2\end{aligned} \tag{S53}$$

**Murein anabolism**

$$\begin{aligned}&-0.11\ \text{ADP} - 0.21\ \text{glucose} - 0.046\ \text{NADP} - 0.21\ \text{NH}_4^+ - 0.17\ \text{O}_2 + 1.00\ \text{murein} \\ &+0.11\ \text{ATP} + 0.18\ \text{CO}_2 + 0.68\ \text{H}_2\text{O} + 0.36\ \text{H}^+ + 0.046\ \text{NADPH}\end{aligned} \tag{S54}$$

Ions are not polymerized into macromolecules, but the concentrations decrease due to dilution by growth, so their uptake is required for cell growth. The glucose consumption in the ion pool energy catabolism is related to transport costs of the ions. Unintuitively, more ATP is required for the transport of ions than for the production of DNA. However, during DNA production, some ATP could be gained between glucose and the DNA precursors, via the energy conserving pathways. No ATP can be obtained for ion transport in such a way.

**Ion pool**

$$\begin{aligned} -0.0052\ Ca^{2+} - 0.0052\ Cl^{-} - 2.5 \cdot 10^{-05}\ Co^{2+} - 0.00071\ Cu^{2+} - 0.015\ Fe^{2+} \\ -0.0019\ \text{glucose} - 0.0078\ H^{+} - 0.2\ K^{+} - 0.0087\ Mg^{2+} - 0.00069\ Mn^{2+} \\ -7 \cdot 10^{-06}\ MoO_4^{2-} - 0.013\ NH_4^{+} - 0.00032\ Ni^{2+} - 0.014\ O_2 - 0.004\ PO_4^{3-} \\ -0.0043\ SO_4^{2-} - 0.00034\ Zn^{2+} + 1.00\ \text{ion pool} + 0.012\ CO_2 + 0.016\ H_2O \end{aligned} \tag{S55}$$

**Ion pool energy catabolism**

$$-0.047\ ADP - 0.0019\ \text{glucose} - 0.012\ O_2 + 0.047\ ATP + 0.012\ CO_2 + 0.012\ H_2O \tag{S56}$$

**Ion pool anabolism**

$$\begin{aligned} -0.047\ ATP - 0.0052\ Ca^{2+} - 0.0052\ Cl^{-} - 2.5 \cdot 10^{-05}\ Co^{2+} - 0.00071\ Cu^{2+} \\ -0.015\ Fe^{2+} - 0.0078\ H^{+} - 0.2\ K^{+} - 0.0087\ Mg^{2+} - 0.00069\ Mn^{2+} \\ -7 \cdot 10^{-06}\ MoO_4^{2-} - 0.013\ NH_4^{+} - 0.00032\ Ni^{2+} - 0.002\ O_2 - 0.004\ PO_4^{3-} \\ -0.0043\ SO_4^{2-} - 0.00034\ Zn^{2+} + 1.00\ \text{ion pool} + 0.047\ ADP + 0.0039\ H_2O \end{aligned} \tag{S57}$$

Similar to ions, cofactors are also diluted by growth and thus need to be produced. Cofactor synthesis consumes 2.7% of the total glucose. For the production of the cofactors, small amounts of ATP and NADPH are required.

**Cofactor pool**

$$\begin{aligned} -2 \cdot 10^{-06}\ \text{biotin} - 0.0015\ Fe^{2+} - 0.027\ \text{glucose} - 0.032\ NH_4^{+} - 0.035\ O_2 \\ -0.0083\ PO_4^{3-} - 0.0022\ SO_4^{2-} + 1.00\ \text{cofactor pool} + 0.034\ CO_2 + 0.14\ H_2O + 0.025\ H^{+} \end{aligned} \tag{S58}$$

**Cofactor pool energy catabolism**

$$\begin{aligned} -0.032\ ADP - 0.0046\ \text{glucose} - 0.0036\ H_2O - 0.031\ NADP - 0.012\ O_2 + 0.032\ ATP \\ +0.027\ CO_2 + 0.062\ H^{+} + 0.031\ NADPH \end{aligned} \tag{S59}$$

**Cofactor pool anabolism**

$$\begin{aligned} -0.032\ ATP - 2 \cdot 10^{-06}\ \text{biotin} - 0.0015\ Fe^{2+} - 0.022\ \text{glucose} - 0.037\ H^{+} \\ -0.031\ NADPH - 0.032\ NH_4^{+} - 0.023\ O_2 - 0.0083\ PO_4^{3-} - 0.0022\ SO_4^{2-} + 1.00\ \text{cofactor pool} \\ +0.032\ ADP + 0.007\ CO_2 + 0.14\ H_2O + 0.031\ NADP \end{aligned} \tag{S60}$$

Growth associated maintenance is ATP hydrolysis for non-metabolic purposes. 16.12 mmolATP/gDW is required for non-metabolic purposes in *E. coli*. This is 56.9% of the ATP required from energy catabolism.

**Growth associated maintenance**

$$-16.12\ ADP - 0.67\ \text{glucose} - 4.03\ O_2 + 16.12\ ATP + 4.03\ CO_2 + 4.03\ H_2O \tag{S61}$$

All MCEQs of macromolecules sum up to the MCEQ of growth and all energy catabolism and anabolism MCEQs of the macromolecules sum to the total energy catabolism and anabolism, respectively.

#### B.3 Assessing the variability of the energetic properties within the optimal solution space

The optimal solution to an FBA is not unique. To assess the robustness of the results of the separation of catabolism and anabolism to this variability, we sampled some alternative optimal solutions of *E. coli* growing aerobically on glucose and considered their energetic properties (Table S3). To sample the alternative optimal solutions, we performed flux variability analysis (FVA) on all fluxes, while keeping the biomass flux and the glucose flux fixed to their optimal value. FVA was performed in a model where the reactions were not split into forward and reverse reactions, to prevent variability of 1000 in each of these reactions, where any flux is canceled out by the reverse flux. No exchange fluxes had variability, which indicates that the macrochemical equation is constant for all variability. Next, we maximized a random flux with value 0 and a non-zero variability. Consequently, another flux became inactive. We set the lower and upper bound of the flux(es) that became inactive to 0 to deactivate it. There were duplicate sets of fluxes to deactivate. For 83 reactions that had variability, there were 11 unique sets of fluxes to deactivate. Then, we used this model to perform the disentanglement between catabolism and anabolism. In short, there was no variability in the energetic properties when choosing an alternative optimal solution (Table S3).

#### B.4 Freedom in energy carrier choice for energy catabolism

In most cases, ATP is exchanged between catabolism and anabolism. However, we have shown that ATP is not always exchanged, e.g. in the case *C. ljungdahlii* growing on $CO_2$ and $H_2$, no ATP is exchanged. However, we have also shown that in the case of *E. coli*, NADH could be exchanged instead of ATP, and we chose ATP. Contrarily, for photosynthetic growth of *Synechocystis*, faced with the choice between ATP and NADH, we chose NADH as exchanged energy carrier. We based this choice on our view of metabolism. If we had chosen NADH in *E. coli*, no ATP would have been exchanged at all. If we had chosen ATP in *Synechocystis*, ATP would have been made from only photons, without any electron donor/acceptor present.

In the method presented here, either ATP or NADH is chosen as energy carrier, and not a combination of the two. It is not possible to determine a unique combination of the two, as they are interchangeable without energy and carbon cost and the exchange of energy between NADH and ATP also occurs in anabolism.

Not only does the choice of energy carrier affect the actual exchange of the energy carrier, but also which and how much electron donor and acceptor is allocated to catabolism and anabolism. This results from the definition of catabolism and anabolism as elementary and electron balanced. Contrary to ATP, NAD(P)H is a redox cofactor. So if a reduced compound is a product of catabolism, less (more) electron acceptor (donor) is required in catabolism to solve the electron balance. The overall macrochemical equation remains fixed, as it resulted from the first optimization of biomass.

Essentially, we highlighted the importance of choice in energy carrier here. In most cases, ATP can be chosen. However, consideration of the specific metabolic pathways of certain organisms is important.

#### B.5 Flux variability when separated on carbon precursors

The precursor set and the relative amounts of precursors produced in precursor catabolism are not unique. Here, we illustrate this with the case of *E. coli* growing aerobically on glucose.

To analyze the possible variability in carbon precursor consumption and production in the anabolic subnetwork, we enumerated the EFMs of the network, with available energy carrier and carbon precursor supplies using EFMTool via CNApy [15]. The network contained 806 EFMs, 506 of which were growing EFMs. Of those EFMs, the variability in energy carrier and carbon precursor consumption and production were considered (Figure S2). There are some metabolites of which the production or consumption does not have any variability (Figure S2D). Some EFMs have high fluxes of carbon precursors, that correlate with each other, for example pep and oxaloacetate (red dots in Figure S2B) and fumarate, pyruvate, ribose-5P and 3P-glycerate (green dots in Figure S2C). The variability in

Table S3: Comparison of metabolic fluxes and deactivated reactions.

| Index | ATP transferred [mmol/gDW/h] | ATP fraction [-] | NADPH transferred [mmol/gDW/h] | Deactivated fluxes |
|---|---|---|---|---|
| 1 | 28.331699 | 0.618230 | 12.178786 | R_FRD2, R_FBA, R_IPDPS, R_ASPO6, R_GLUt2rpp, R_GLUt4pp, R_DUTPDP, R_PFK, R_RNTR4c2, R_PYK3, R_SUCCt1pp |
| 2 | 28.331699 | 0.618230 | 12.178786 | R_FBA, R_IPDPS, R_GLUt2rpp, R_GLUt4pp, R_DUTPDP, R_PFK, R_RNTR4c2, R_PYK3, R_SUCCt1pp |
| 3 | 28.331699 | 0.615894 | 12.178785 | R_DHORD5, R_FRD2, R_FBA, R_IPDPS, R_ASPO6, R_GLUt2rpp, R_GLUt4pp, R_DUTPDP, R_PFK, R_RNTR4c2, R_PYK3, R_SUCCt1pp |
| 4 | 28.331699 | 0.618230 | 12.178785 | R_FBA, R_IPDPS, R_ASPO6, R_GLUt2rpp, R_GLUt4pp, R_PFK, R_PYK3, R_SUCCt1pp |
| 5 | 28.331699 | 0.615894 | 12.178785 | R_FBA, R_ASPO6, R_GLUt2rpp, R_GLUt4pp, R_DUTPDP, R_PFK, R_RNTR4c2, R_PYK3, R_SUCCt1pp |
| 6 | 28.331699 | 0.615894 | 12.178785 | R_FBA, R_IPDPS, R_ASPO6, R_GLUt2rpp, R_GLUt4pp, R_DUTPDP, R_PFK, R_G3PAT161, R_RNTR4c2, R_PYK3, R_SUCCt1pp |
| 7 | 28.331699 | 0.618230 | 12.178786 | R_IPDPS, R_ASPO6, R_GLUt2rpp, R_GLUt4pp, R_DUTPDP, R_RNTR4c2, R_PYK3, R_SUCCt1pp |
| 8 | 28.331699 | 0.618230 | 12.178786 | R_FBA, R_IPDPS, R_GART, R_ASPO6, R_GLUt2rpp, R_GLUt4pp, R_DUTPDP, R_PFK, R_RNTR4c2, R_PYK3, R_SUCCt1pp |
| 9 | 28.331699 | 0.615894 | 12.178785 | R_FBA, R_IPDPS, R_ASPO6, R_GLUt2rpp, R_GLUt4pp, R_DUTPDP, R_PFK, R_RNTR4c2, R_PYK3, R_SUCCt1pp |
| 10 | 28.331699 | 0.618230 | 12.178785 | R_FBA, R_IPDPS, R_GLUt2rpp, R_GLUt4pp, R_DUTPDP, R_QMO3, R_PFK, R_RNTR4c2, R_PYK3, R_SUCCt1pp |
| 11 | 28.331699 | 0.615894 | 12.178785 | R_FBA, R_IPDPS, R_ASPO6, R_GLUt2rpp, R_GLUt4pp, R_DUTPDP, R_PFK, R_NADH16pp, R_RNTR4c2, R_PYK3, R_SUCCt1pp |

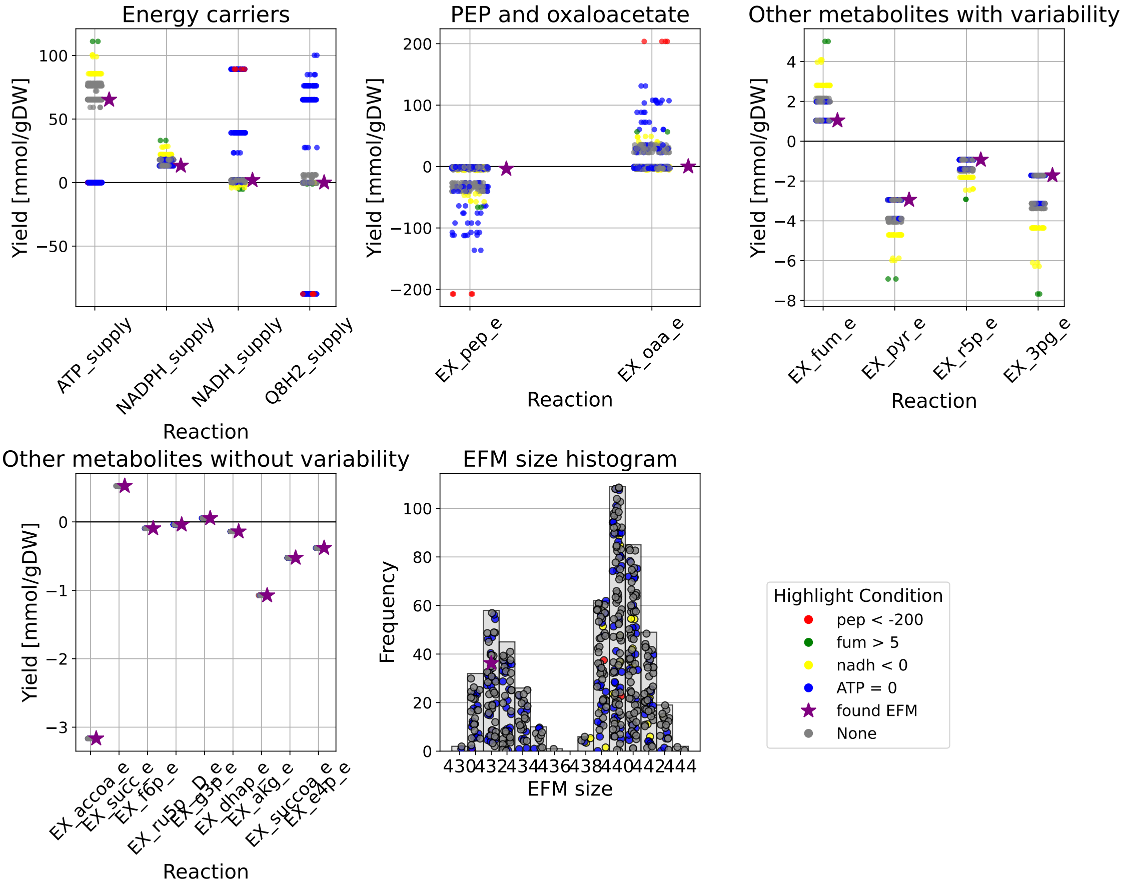

Figure S2: Yields of energy carrier (A) and carbon precursor (B, C and D) consumption in different EFMs. Each dot corresponds to an EFM. (E) The number of reactions in the EFMs.

energy carriers is also correlated: NADH (NAD ¡ 0, yellow dots) production in anabolism correlates with high ATP and NADPH consumption, as well as higher consumption of the variable metabolites. In addition, no consumption (or production) of ATP (blue dots) correlates with higher NADH and Q8H2 consumption.

To conclude, part of the variability can be easily explained by precursors and energy carriers being converted into one another. There is still some variability left (grey dots) that might be due to the use of alternative pathways for biomass production. We choose to move on with one of the solutions that was found, but do not draw any conclusions based on carbon precursor consumption that is potentially variable.

#### B.6 Producing macromolecules together is more efficient than separately

Adding up the flux distributions for the production of each macromolecule separately does not add up to the originally found flux distribution when optimizing for biomass. More glucose is consumed when optimizing the macromolecule productions separately (Figure S3). This is due to the fact that some internal metabolites can be shared when simultaneously optimizing production of all macromolecules (i.e. optimizing biomass production), that have to be balanced each time when optimizing the production of each macromolecule separately. An example of this is that lipid production has ATP as a byproduct (see Section **??**), that can be used directly for another macromolecule in the biomass optimization, but has to be hydrolyzed to ADP when optimizing solely for lipid. As all optimizations were done in the active network, any ATP hydrolysis step might have accompanying carbon costs.

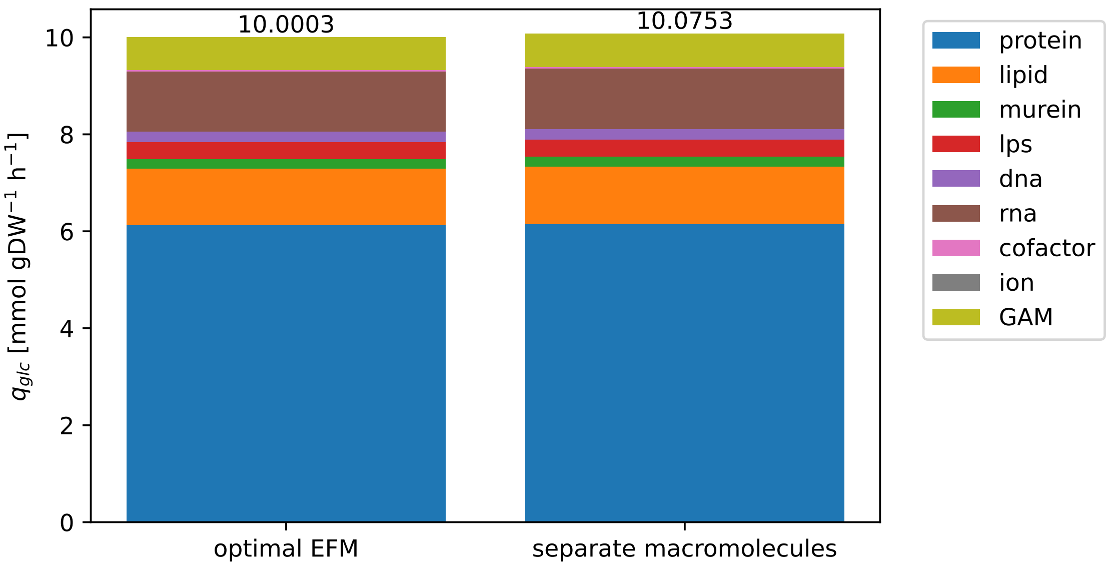

Figure S3: Glucose consumption (mmol gDW$^{-1}$ h$^{-1}$) for macromolecules separated from the biomass-optimal EFM (left bar) and the sum for each macromolecule separately (right bar).

#### B.7 No single determinant exists for the role of energy catabolism

To determine whether the energy-related parameters could be explained by e.g. properties of the substrate, or overall biomass yield on the energy source, we plotted the relationships against each other (Figure S4 and S5). Logically, yield on the substrate, per carbon atom in the energy source and per electron in the carbon source correlate with each other. The only energy-related parameters are the amount of ATP transferred and the fraction of ATP produced during precursor synthesis, which is a trivial relationship. Thus, we conclude that precursor catabolism must be taken into account, as no conclusions can be drawn from dissection based on energy alone.

#### B.8 The 12 classic precursors for growth do not solely support growth

According to Neidhardt (1990) [16], any cell is able to make itself from the following 12 metabolites: 3P-glycerate, acetyl-CoA, erythrose-4P, fructose-6P, glyceraldehyde-3P, glucose-6P, oxaloacetate, phosphoenolpyruvate, pyruvate, ribose-5P, α-ketoglutarate and succinyl-CoA. We aim to test this hypothesis using a genome scale model of *E. coli* [4]. First, the active model for optimal growth on glucose and $O_2$ was obtained (see Section A.1), to which exchange and transport reactions are added for the 12 precursors, according to for example equations S14-S16. Next, ATP, NADH and NADPH supply were added, according to equations S5 and S6. Then, to disable the model from using pathways in central carbon metabolism to convert one carbon precursor into the other, the following reactions were deactivated: GAPD_fwd, PYK, PGK_rev, G6PDH2r_fwd, GND, PGL, PGI_fwd, FBA_fwd, PFK, TPI_fwd, CS, FUM_fwd, TKT1_fwd, TKT2_fwd, PDH, AKGDH, ACONTa_fwd, ACONTb_fwd, ICDHyr_fwd, SUCDi, SOCOAS_rev, MDH_fwd, GLCptspp, TALA_fwd, PGM_rev. Then, the lower and upper bound of the BOF were set to 1.0 and the BOF was optimized (to find any feasible flux distribution). There was no feasible flux distribution.

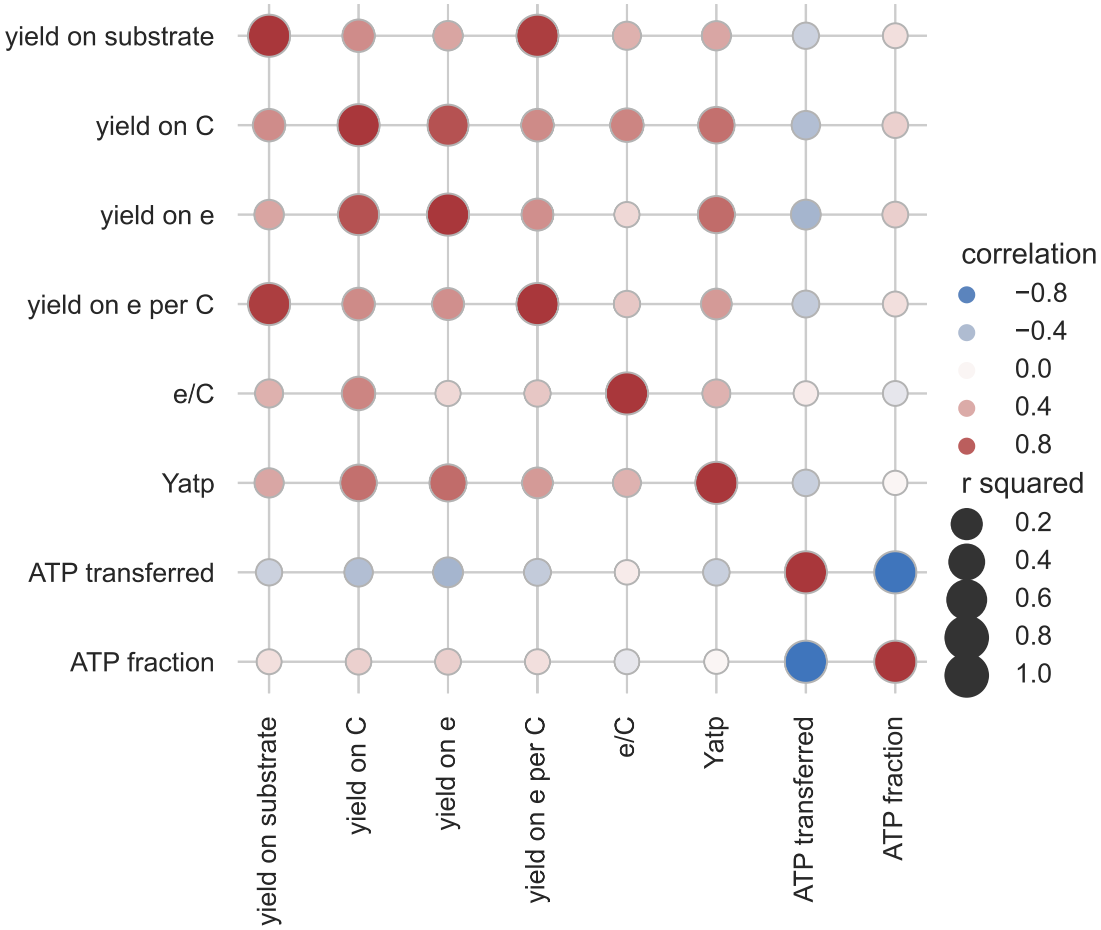

Figure S4: Correlation of components of the macrochemical equation with each other and the fraction of ATP produced in anabolism. Units of the parameters plotted are: yield on substrate [gDW $mmol_S^{-1}$], yield on C [gDW $mCmol_S^{-1}$], yield on e [gDW $memol_S^{-1}$], yield on e/C [gDW $mCmol_S$ $memol_S^{-1}$], e/C [$emol_S$ $Cmol_S^{-1}$], $Y_{ATP}$ [gDW $mol_{ATP}^{-1}$], ATP transferred [$mmol_{ATP}$ $gDW^{-1}$ $h^{-1}$] and ATP fraction [-]. Genome scale models used for these simulations are: iHN637 [11] (*C. ljungdahlii*), iAF987 [12] (*G. metallireducens*), iMG746 [13] (*M. barkeri*), Yeast9 [6] (*S. cerevisiae*), iJN1462 [14] (*P. putida*), iML1515 [4] (*E. coli*) and iSynCJ816 [10] (*Synechocystis*).

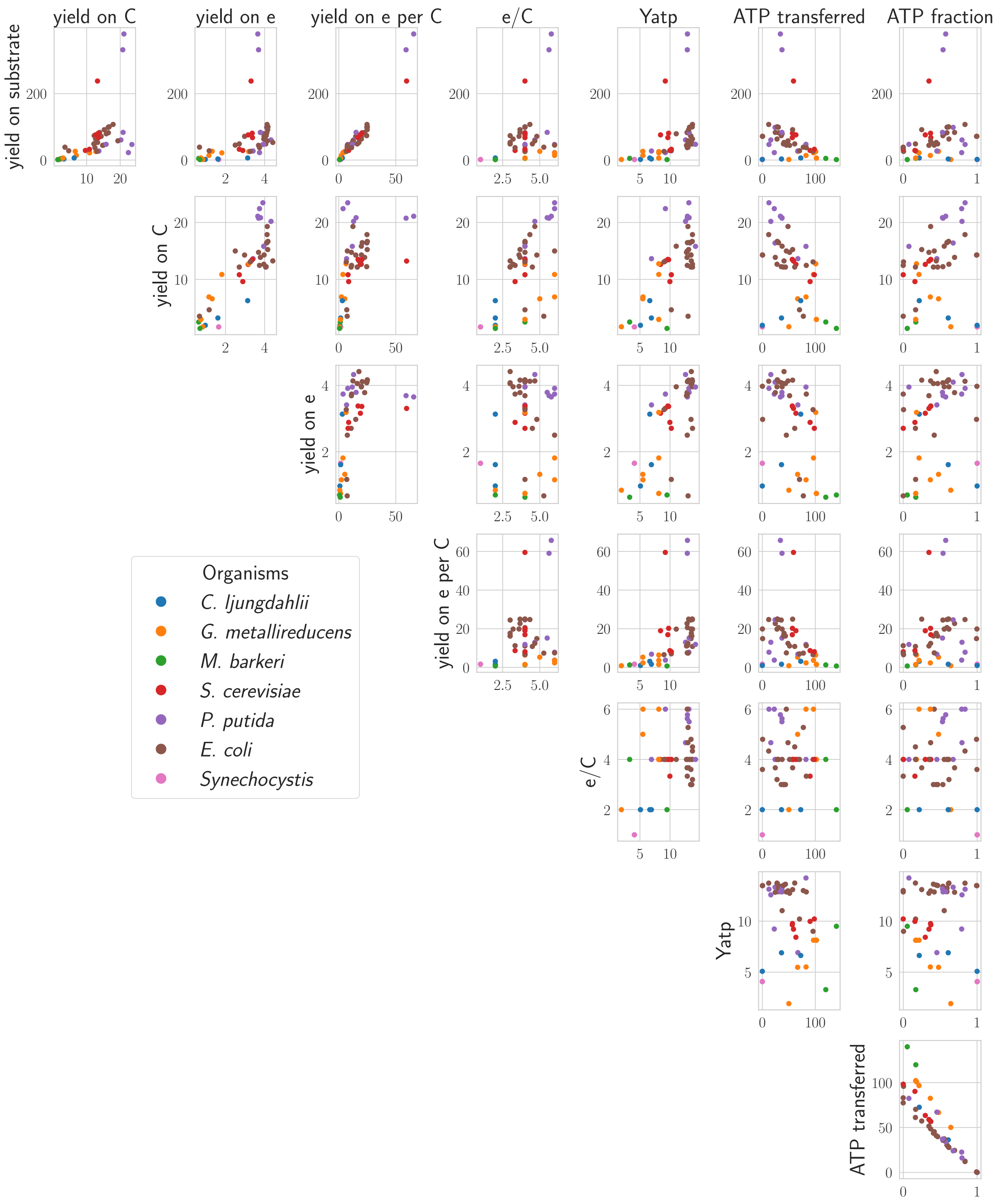

Figure S5: Correlations of biomass yield on substrate [gDW mmol$_{ES}$$^{-1}$], biomass yield per carbon [gDW mCmol$_{ES}$$^{-1}$], biomass yield per electron [gDW memol$_{ES}$$^{-1}$], growth per electron content of the energy source [gDW mCmol$_{ES}$$^{-1}$ memol$_{ES}$$^{-1}$], electron content per carbon in the energy source [e$^{-}$ C$^{-1}$], ATP yield [gDW mol$_{ATP}$], ATP transferred from catabolism to anabolism [mmol gDW$^{-1}$ h$^{-1}$] and fraction of ATP produced in anabolism [-] against each other.

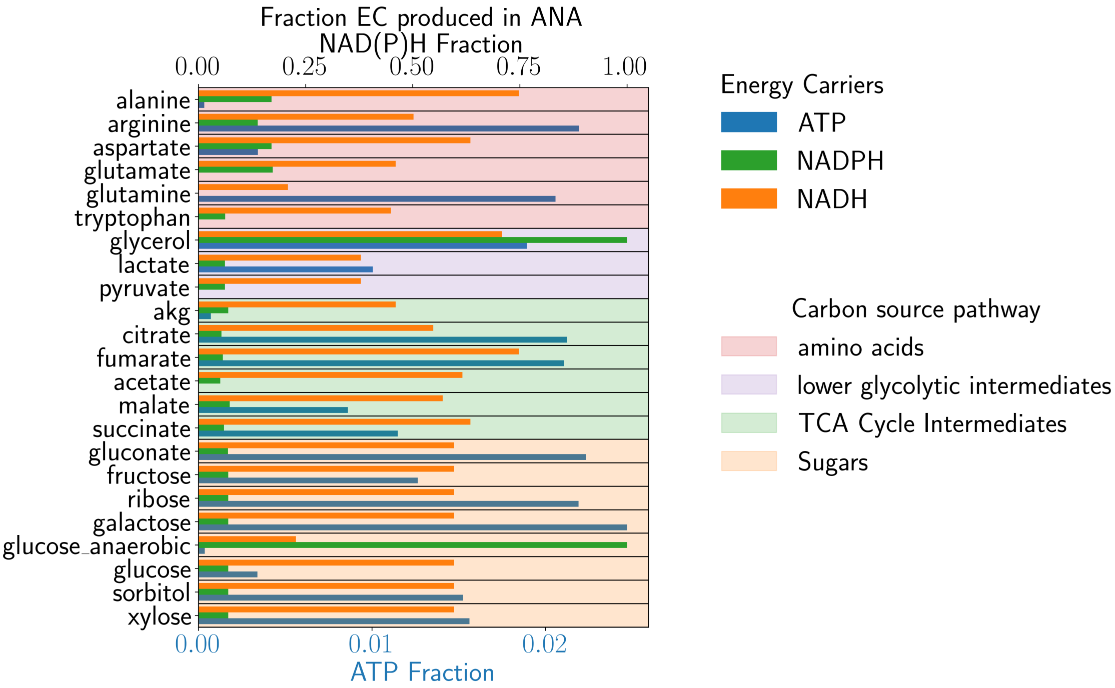

Figure S6: Caption

[16] Frederick Carl Neidhardt, John L Ingraham, and Moselio Schaechter. Physiology of the bacterial cell: a molecular approach. 1990.
